# A single-cell transcriptomic atlas maps cerebellar astrocyte diversity and uncovers the transcriptional code underlying their maturation trajectories

**DOI:** 10.1101/2025.07.17.665323

**Authors:** Valentina Cerrato, Giacomo Turrini, Ilaria Vitali, Bilian Xiong, Laura Solanelles-Farré, Andrea Lopes, Elia Magrinelli, Riccardo Bocchi, Magdalena Götz, Judith Fischer-Sternjak, Enrica Boda, Annalisa Buffo, Ludovic Telley

## Abstract

Astrocytes are increasingly recognized as key regulators of neural circuit development and function, with mounting evidence revealing substantial heterogeneity within and across brain regions. Yet, the full extent of this diversity and its developmental mechanisms remain poorly understood. To address this, we leveraged the uniqueness of the mouse cerebellum, which hosts well-defined astrocyte types and established progenitor pools. Through complementary multi-modal omic approaches, including single-cell RNA sequencing, spatial transcriptomics, trajectory inference, clonal lineage reconstruction, and gene expression and regulatory network analyses, we systematically dissected the molecular diversity and ontogenesis of cerebellar astrocytes. We identified known types and uncovered new subtypes with functional specialization, inferring their developmental trajectories from multiple embryonic niches and postnatal progenitor sources with fate divergence, convergence, and restriction. We further predicted a hierarchical transcriptional regulator code governing this diversification, operating at multiple levels: distinct regulatory modules i) reflect embryonic regionalization and lineage; ii) determine broad astroglial identity; specify iii) Bergmann versus non-Bergmann fates; and guide iv) astrocyte type and v) subtype acquisition. Our findings map and temporally organize transcriptional programs that capture key determinants of astrocyte fate, integrating them along defined trajectories toward diverse astrocyte identities. This high-resolution framework for cerebellar glial diversification offers a model to be challenged across other brain regions.

## Introduction

Over recent decades, astrocytes have emerged as central players in neural circuit development, function, and plasticity, shifting the traditional notion of their role from merely supportive elements to active participants in CNS physiology^1,2^. Accumulating evidence underscores the substantial molecular, morphological, and functional heterogeneity among astrocytes, which aligns with their distinct regional identities, developmental origins, and specialized functions^3–9^.

Astrocyte diversity can be remarkably profound, even within a single anatomical region. For instance, in the cerebral cortex, astrocytes demonstrate transcriptional heterogeneity that partially reflects the cortical laminar structure^10,11^. Similarly, distinct astrocyte subtypes have been identified within the layered architecture of the hippocampal dentate gyrus^12^.

Nevertheless, clearly defining astrocyte types and subtypes within brain regions remains challenging, as subtle morphological and molecular variations frequently blur distinctions between astrocyte populations, complicating precise identity and functional definitions.

Thus, despite substantial progress, achieving a comprehensive characterization of astrocyte heterogeneity, particularly within individual brain regions, and elucidating the developmental mechanisms underlying this diversity, remain open goals.

To address these challenges, we leveraged the robust framework provided by the mouse cerebellum, distinguished by well-defined astrocyte types, with stereotyped morphologies, spatial allocations^13^ (Fig. 1d) and established astroglial progenitor lineages^14–16^ and by spatially segregated functional compartments^17^. This knowledge, integrated with single-cell/single-nucleus RNA sequencing (sc/snRNA-seq), maturational trajectory inference, clonal investigations, differential gene expression, and gene regulatory network analysis, allowed to systematically dissect cerebellar astrocyte heterogeneity and its development with unprecedented efficacy.

**Fig. 1.**
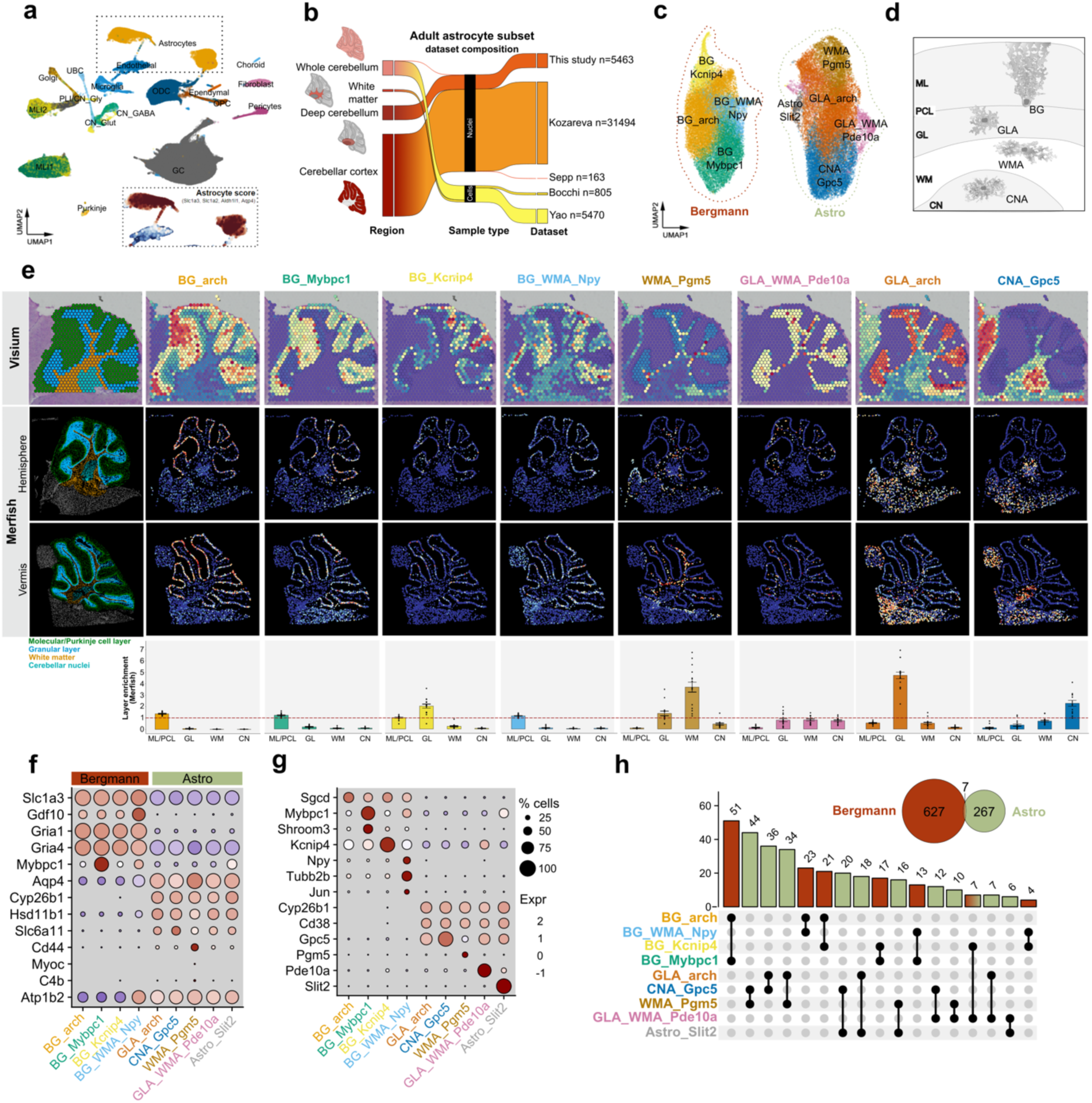
Sc/snRNA-seq resolves main astrocyte types and uncovers new subtypes in the adult mouse cerebellum. a, UMAP plot of the six integrated datasets of the adult mouse cerebellum, with dots colored based on cell types. The inset shows the cells with a high astrocyte score, defined by the combinatorial expression of *Slc1a3*, *Slc1a2*, *Aldh1l1* and *Aqp4.* **b,** Flow diagram showing the dataset composition (i.e. sampled cerebellar region, type of sample, name of the original dataset and number of cells/nuclei per dataset) of the astrocyte subset. n= number of astrocytes. **c,** UMAP plot of the astrocyte subset. The dotted lines surround the two broad groups corresponding essentially to Bergmann glia (Bergman) and the other non-Bergmann glia astrocytes (Astro). **d,** Schematic representation of the main astrocyte types of the cerebellum and their localization in the cerebellar grey or white matter regions: Bergmann glia (BG) in the Purkinje cell layer (PCL) and molecular layer (ML), grey matter astrocytes (GLA) in the granular layer (GL), white matter astrocytes (WMA) in the white matter (WM), and cerebellar nuclear astrocytes (CNA) in the cerebellar nuclei (CN). **e,** Mapping of the identified astrocyte clusters in adult mouse Visium (first line) or Merfish (second and third line) ST datasets. The first column shows the layer annotation of the sagittal sections analyzed. Bar plots at the bottom show the average layer enrichment predicted in multiple (n=16) sections of the Merfish dataset (+-SEM). Cells mapped in the ML/PCL, GL, WM or CN were annotated as BG, GLA, WMA or CNA, respectively. **f,** Expression of marker genes supposed to be enriched or specific for the distinct astrocyte types (*Slc1a3, Gdf10, Gria1, Gria4* for BG; *Aqp4, Cyp26b1, Hsd11b1* for GLA; *Slc6a11* for CNA; *Cd44, Myoc, C4b* for WMA; *Atp1b2* for all astrocytes except BG) or subtypes (*Mybpc1* for Bergmann_2 cluster in Kozareva et al.^18^). **g,** Expression of the top DEGs (avgLog2FC > 1) of each of the identified astrocyte clusters. **h,** Pairwise cumulative comparisons across the DEGs of each cluster. Note that most overlaps are between clusters belonging either to the Bergmann or to the Astro groups, with the only exception of BG_*Kcnip4* and GLA_WMA_*Pde10a*, that share 7 genes (intersection highlighted in the Venn diagram). BG, Bergmann glia; GLA, granular layer astrocytes; WMA, white matter astrocytes; CNA, cerebellar nuclei astrocytes; ML, molecular layer; PCL, Purkinje cell layer.

Building upon this knowledge, this work identifies so far unrecognized astrocyte subtypes, revealing unique spatial distributions and functional specializations. We delineate the transcriptional regulator code orchestrating cerebellar astrocyte heterogeneity and show that this diversity arises from both canonical and non-canonical intra-and extra-cerebellar astrogliogenic niches, involving complex patterns of cell fate divergence, convergence, and restricted lineage paths from multiple progenitor sources.

## Results

### Transcriptional heterogeneity resolves main adult astrocyte types and uncovers new subtypes

To address the intra-regional diversity of cerebellar astrocytes, we first looked at existing sc/snRNA-seq datasets of the adult mouse cerebellum^18–21^. However, we found that they provided limited coverage of non-cortical cerebellar astrocytes, restricting a comprehensive investigation of these cell populations (Extended Data Fig. 1a). To overcome this limitation, we generated new sc/snRNA-seq data through manual microdissection of deep cerebellar regions, specifically of the cerebellar nuclei (this work, Supplementary Table 1) or deep white matter^22^. These new data were integrated *in silico* with the existing published datasets, resulting, after quality control and filtering (see Methods), in a combined total of 493,505 high quality nuclei/cells (hereafter referred to as “cells”) with a total of 22,335 genes detected (Fig. 1a).

Following unsupervised clustering in Seurat V5 and based on the expression of cell type-specific molecular markers, we assigned these cells to 18 distinct cell types (Fig. 1a), comprising 7 classes and 10 subclasses^18,21^ (Extended Data Fig. 1b). Cells with a high (>1) astrocyte score, determined by the combined expression of established astrocyte-specific markers^7,9^ (*Slc1a3*, *Slc1a2*, *Aldh1l1* and *Aqp4*), were then subsetted for further analyses, resulting in an integrated astrocyte dataset comprising a total of 43,395 astrocytes and 19,914 genes detected. This provided comprehensive coverage and high resolution across the entire cerebellum (Fig. 1b). Astrocytes were distributed into 9 clusters within the UMAP embedding, forming two segregated groups (Fig 1c). Mapping to two publicly available adult mouse spatial transcriptomics (ST) datasets^20^ allowed to infer that the two broad groups were composed of distinct astrocyte types of the grey and white matter, matching the canonical classification of cerebellar astrocytes, comprising Bergmann glia (BG) and velate granular layer astrocytes (GLA) in the cortex, white matter astrocytes (WMA), and cerebellar nuclei astrocytes (CNA; Fig. 1d,e). Specifically, one group was predominantly composed of Bergmann glia (Bergmann), and the second group of non-Bergmann astrocytes (Astro). Astrocyte subtypes were also identified, either confined to discrete regions or spanning multiple regions (Fig. 1e).

When looking for the genes at the core of molecular diversity across clusters, the sole expression levels of most of the markers known to be enriched or specific for certain cerebellar astrocyte types^23–26^ were insufficient on their own to annotate all clusters and, rather, only distinguished Bergmann from Astro (Fig. 1c,f). Finer cluster discrimination was instead achieved with the most specific differentially expressed genes (DEGs) of each cluster (Fig.1g and Supplementary Table 2). Hence, final cluster annotation (Fig. 1c) was obtained by combining regional distribution predictions with the most specific DEGs of each cluster. Two clusters predicted to map to the PCL or GL did not exhibit any enriched DEGs, denoting a reference transcriptomic profile, and were therefore annotated as BG_*arch* (i.e. “archetype”) and GLA_*arch*.

This classification revealed 6 clusters with predominant localization in a single region (BG_*arch*, BG_*Mybpc1*, BG_*Kcnip4*, WMA_*Pgm5*, GLA_*arch* and CNA_*Gpc5;* Fig. 1e). Among the clusters mapping in the ML/PCL, BG_*arch* and BG_*Mybpc1* had a strikingly complementary distribution in the anterior or posterior lobules, respectively (Fig. 1e, Extended Data Fig. 2d,e). Moreover, these 2 clusters in Merfish predictions showed low spatial and transcriptional overlap within each other and with BG_*Kcnip4* (Extended Data Fig. 2h), in line with these 3 clusters representing distinct Bergmann subtypes.

In contrast, 2 clusters (BG_WMA_*Npy* and GLA_WMA_*Pde10a*) spanned across multiple regions and may therefore represent new subtypes of astrocytes with similar transcriptional features but diverse spatial allocations. In the Visium dataset, BG_WMA_*Npy* appeared as an hybrid cluster, enriched in both the PCL/ML and WM (Fig. 1e). This enrichment varied depending on the cell source: cells collected from the whole cerebellum or the cerebellar cortex^18–20^ mapped to the PCL/ML, whereas cells sourced from the deep cerebellum (this work, Bocchi et al.^22^) mapped to the WM (Extended Data Fig. 2a,b). The cluster hybrid nature was further confirmed by its overlapping signature with the cerebellar WMA cluster “Cer_2” of Bocchi et al.^22^ (Extended Data Fig. 1c) in conjunction with high expression of typical Bergmann genes (Fig. 1f). The BG_WMA_*Npy* cluster was enriched with gene terms associated with translation and protein folding, processes typically upregulated in neural progenitors and stem cells^27,28^, and exhibited specific enrichment of genes expressed in NSCs and radial glial cells (Fig. 1g, Extended Data Fig. 1d and Supplementary Table 3), in line with the display of an immature profile^22^.

GLA_WMA_*Pde10a* cells were enriched in both the GL and WM. This cluster shared a few DEGs with BG_*Kcnip4*, a unique case of overlap between Bergmann and Astro groups (Fig. 1h). Both these subtypes also displayed a high glutamate release score, defined by the expression of *Syt1*, *Snap25* and *Slc17a7*, and shared many DEGs with the recently uncovered specialized astrocyte subtype that mediates glutamatergic gliotransmission^29^ (Extended Data Fig. 1e). Accordingly, GO analysis highlighted for GLA_WMA_*Pde10a* an enrichment of terms related to synaptic vesicle exocytosis and synaptic transmission (Extended Data Fig 1f, Supplementary Table 3). These clusters may therefore represent functionally specialized vesicular-releasing subtypes particularly involved in gliotransmission and localized in distinct grey and white matter cerebellar regions.

Of note, the reported analysis showed 3 clusters containing WMA, suggesting distinct subtypes with complementary enrichment patterns in the lobular (WMA_*Pgm5* and WMA_*Pde10a*) or deep WM (WMA_*Npy*; Fig. 1e, Extended Data Fig. 2b). Additionally, we identified a small *Slit2*-expressing cluster exhibiting a salt and pepper pattern (Extended Data Fig. 2c) with apparent enrichment in the posterior lobules of the vermis (IX/X) (Extended Data Fig. 2d,e). Slit2 expression may reflect a reactive-like state^30^ in non-Bergmann glial cells.

Cross-comparisons showed significant transcriptional overlap within the Bergmann and Astro groups, but not between them (Fig. 1h). GO analysis supported this divergence, confirming Bergmann glia main specialization in maintaining cerebellar structural integrity, regulating glutamatergic inputs, and relying on Shh signaling to sustain their molecular identity^24,31^. Conversely, this analysis uncovered novel potential roles for non-Bergmann Astro, including involvement in energy homeostasis via regulation of blood supply, glucose and glycogen metabolism, and neuroprotection (Extended Data Fig. 1g, Supplementary Table 4).

Overall, these data point to spatially segregated astrocyte subtypes, some of which exhibit features suggestive of functional specializations. They also indicate remarkable functional divergence between Bergmann and non-Bergmann astrocytes.

### Transcriptional heterogeneity defines postnatal astroglial-like progenitor pools transitioning into fully mature astrocyte subtypes

To elucidate the developmental processes underlying the remarkable heterogeneity of cerebellar astrocytes, we generated an integrated multi-modal sc/snRNAseq dataset of the postnatal cerebellum. This dataset encompasses 28 distinct experiments conducted across developmental stages from postnatal day (P) 0 to P30 (P0, P4–P5, P7, P10–P11, P14–P16, P23, P30) and integrates diverse information, partly exploited in this study, including time of birth, clonal relationships, antero-posterior (AP) and medio-lateral (ML) spatial resolution (Extended Data Fig. 3a, Supplementary Table 1). Cells collected at biologically similar time points (P0-P5: early postnatal development; P7-P11: circuit assembly; and P14-P30: final maturation and refinement) were integrated *in silico.* Following quality control and filtering steps (see Methods), the dataset retained 126,143 high quality cells (P0-P5 = 85,443; P7-P11 = 19,105; P14-P30 = 21,645), with a total of 20,226 genes detected across all stages. Following unsupervised clustering, a deep neural network-based prediction model^29^ combined with manual annotation based on known marker genes were used to classify the cells (Fig. 2a; Extended Data Fig. 3b). Comparison with publicly available datasets of the postnatal mouse cerebellum^19,32,33^ demonstrated the unique resolution of this dataset, capturing the full repertoire of cerebellar cell types with exceptional detail (Extended Data Fig. 3c).

**Fig. 2.**
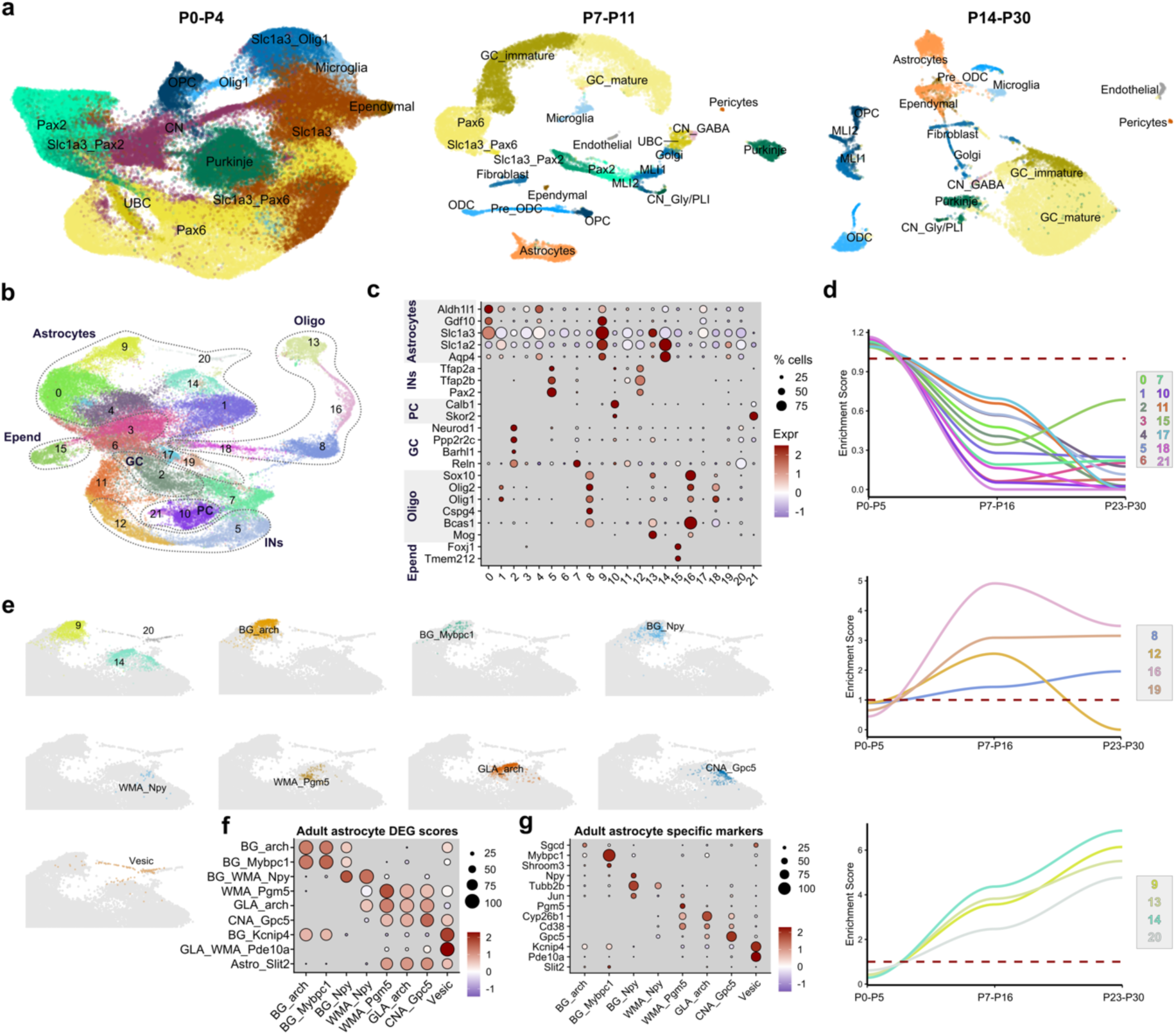
Transcriptomic profiling of the postnatal mouse cerebellum to reveal astrocyte subtypes maturation. a, UMAP plots of the integrated P0-P4, P7-P11 and P14-P30 datasets of the postnatal mouse cerebellum, with dots colored by cell cell types and maturational states. **b,** UMAP plot of the postnatal subset of *Slc1a3-*expressing cells, with dotted lines outlining clusters corresponding to distinct cell lineages. Astrocytes (Astro), GABAergic interneurons (INs), Purkinje cells (PC), glutamatergic granule cells (GC), oligodendrocytes (Oligo), and ependymal cells (Epend). **c,** Expression of lineage-specific markers used to classify cells. **d,** Enrichment scores of distinct clusters across developmental stages, grouped based on their enrichment patterns at early (top), middle (middle) or late (bottom) stages. **e,** UMAP insets from panel **b** highlighting clusters 9, 14 and 20 containing the mature astrocyte subtypes. Annotated subtypes within these clusters are shown in separate insets. **f,** Dot plots of the expression scores of adult astrocyte subtype DEGs. **g,** Dot plots of the expression levels of adult subtype-specific markers.

We used this dataset to subset astroglial-like progenitors and maturing astrocytes based on expression of the pan astrocyte and astroglial-like progenitor marker *Slc1a3*^14,16^. *Slc1a3* expression partially overlapped with other lineage-specific markers, including *Pax2* (GABAergic interneurons; “Slc1a3_Pax2”), *Pax6* (Glutamatergic neurons, “Slc1a3_Pax6”), and *Olig1* (Oligodendrocytes, “Slc1a3_Olig1”). Furthermore, Slc1a3 expression was also detected in cells of the ependymal, microglial, and fibroblast lineages (Extended Data Fig. 3b,d). For further analyses, the *Slc1a3*-expressing clusters were filtered, excluding microglia and fibroblasts due to their non-neural origin, and subsequently re-integrated (Extended Data Fig. 3d). In addition, the entire oligodendrocyte and interneuron lineages, also comprising *Slc1a3*-negative cells, were incorporated into this subset in view to ensure robust internal validation of cell maturation trajectories (see Methods and Extended data Fig.5).

Unsupervised clustering partitioned the *Slc1a3*-expressing subset into 22 distinct clusters, most of which could be grouped into separate lineages based on reference genes (Fig. 2b,c). Specifically, clusters 0, 1, 4, 9, 14, 17 and 19 expressed cerebellar astrocyte-typical genes; clusters 5 and 12 belonged to the GABAergic interneuron lineage; clusters 10 and 21 expressed Purkinje cells (PC)-typical genes; cluster 2 displayed granule cell (GC)-specific features; clusters 8, 13, and 16 represented distinct stages of oligodendrocyte differentiation; and cluster 15 corresponded to ependymal cells. These results align with the presence of *Slc1a3*+ ancestors for astrocytes, interneurons, oligodendrocytes, GCs, and ependymal cells^14,16,34^ and suggest that some immature PCs retain Slc1a3 mRNA up to P5.

Within this subset, cells were distributed according to their age of collection, forming a continuum of cell states ranging from progenitors to fully mature cells (Extended Data Fig. 4a).

Distinct clusters exhibited diverse enrichment patterns across developmental ages: some clusters, likely representing progenitors and including cycling cells, were enriched at neonatal stages but became underrepresented at later stages (Fig. 2b, top panel; Extended Data Fig. 5a). Other clusters, likely corresponding to intermediate progenitor states, exhibited increased enrichment beginning from middle stages and typically plateaued thereafter (Fig. 2b, middle panel). Additional clusters, reflecting maturing cell states, demonstrated a gradual increase in enrichment, peaking at later developmental stages (Fig. 2c, bottom panel).

Inspection of astrocyte-specific markers and their stage-specific distribution (Fig. 2c; Extended Data Fig. 4b,c) classified clusters 9, 14, and 20 as the most mature astrocyte states. Gene expression patterns and spatial mapping confirmed the presence of astrocyte subtypes previously defined in the adult dataset also in these postnatal samples (Fig. 2e,f,g, Extended Data Fig. 4d). Cluster 9 encompassed most terminal differentiated BG subtypes, including BG_*arch*, BG_*Mybpc1*, and BG_*Npy*. Cluster 14 included WMA_*Npy*, WMA_*Pgm5*, GLA_*arch*, and CNA_*Gpc5*. Notably, the analysis of this dataset discriminated between *Npy*-expressing BG and WMA, which were instead merged into a single cluster in the adult dataset. Conversely, a few cells in cluster 20 exhibited molecular profiles shared between BG_*Knip4* and GLA_WMA_*Pde10a*, with a predicted spatial distribution spanning the PCL, GL, and WM. Due to their vesicular-releasing profile, we hereafter refer to this population as “Vesic” (Extended Data Fig. 4d).

Collectively, this postnatal dataset offers a comprehensive representation of both astroglial-like progenitor heterogeneity and mature astrocyte subtypes, making it well-suited for investigating astrocyte maturation trajectories.

### Inferred maturation trajectories identify postnatal multipotent and committed astroglial-like progenitors

To reconstruct the maturation trajectories of the astrocyte subtypes extending back to perinatal progenitor states, we combined URD and two CellRank kernels, thereby integrating three distinct computational approaches to infer cellular dynamics from sc/snRNAseq data, based on similarities in transcriptional profiles and their distribution over time^35,36^ (Fig. 3a). Outputs were integrated into a consensus heatmap by averaging each clusterwise logit-transformed probability of transitioning into a defined transcriptional state (Fig. 3a, Supplementary Table 5). The predictive power of this strategy was validated by using data from the oligodendrocyte and GABAergic interneuron cell classes (Methods and Extended Data Fig. 5).

**Fig. 3.**
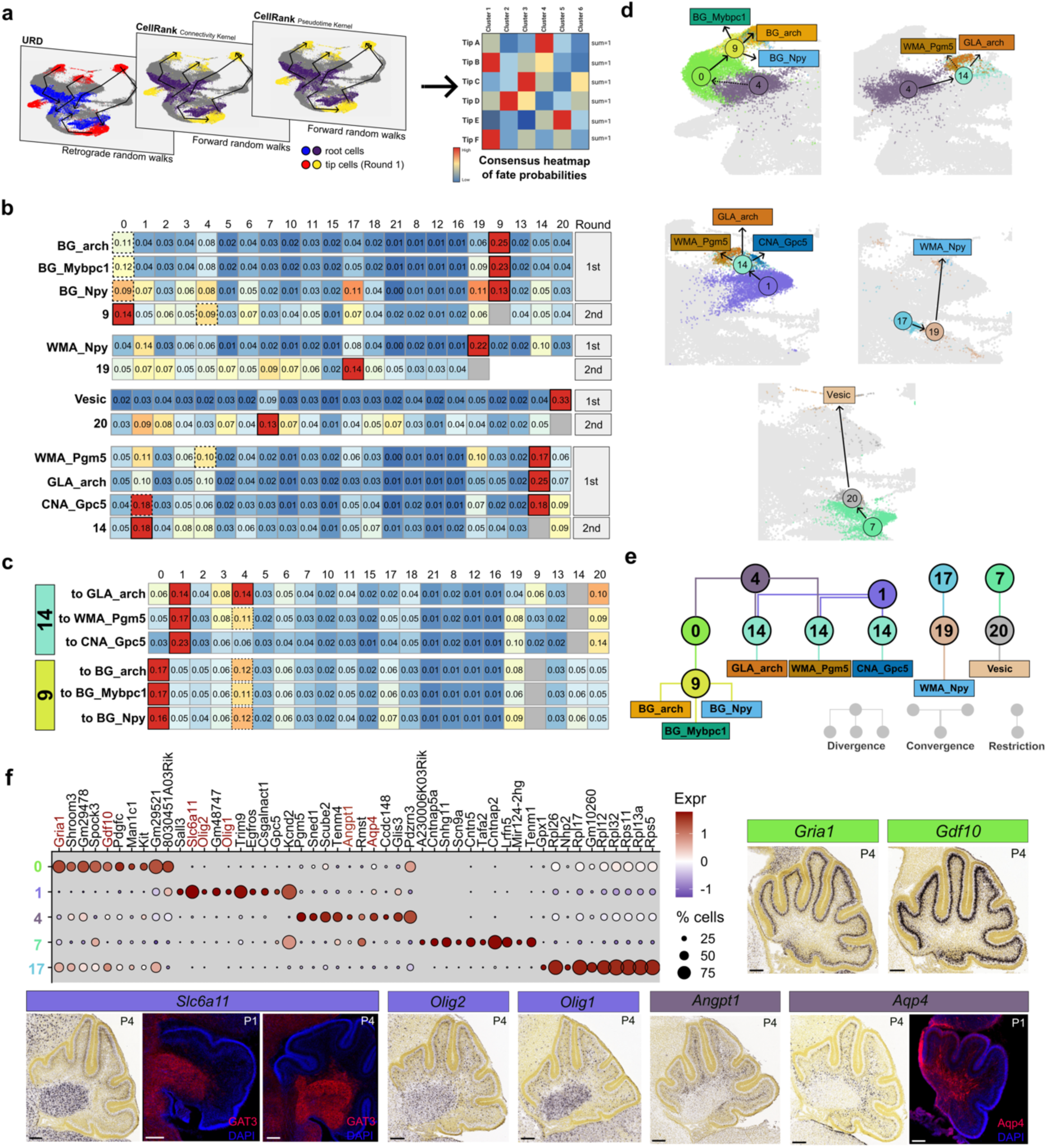
Inferring astrocyte maturation trajectories through a combined computational approach. a, Schematic representation of the *in silico* approach used to predict maturation trajectories, integrating URD^35^ (retrograde random walks) with two distinct CellRank^36^ kernels (forward random walks). Representative plots display the root and tip cells of the 1st rounds of random walks (i.e. from or toward the most differentiated cells), color-coded based on the algorithm used. The clusterwise logit-transformed probabilities from each of the three simulations are averaged to generate a consensus heatmap. **b,** Consensus heatmaps of random walks predicting astrocyte maturation trajectories, showing for each cluster (columns) the probability of differentiating into a defined terminal (1st round) or intermediate (2nd round) fate. Solid and dotted black lines indicate the clusters with the highest or second-highest probability in at least two out of three approaches, respectively. Heatmap colors are scaled independently for each row. **c,** Consensus heatmap of the random walks performed using as tips the cells in cluster 14 (top) or cluster 9 (bottom) with the highest probability of differentiating into GLA_*arch*, WMA_*Pgm5*, CNA_*Gpc5*, or BG_*arch*, BG_*Mybpc1*, BG_*Npy*. Notably, cluster 4 shows the second-highest probability of differentiating into WMA_Pgm5 in both this additional round and the 1st round of random walks (panel **b**, ninth row of the heatmap). **d,** Schematic reconstruction of the differentiation trajectories of each mature astrocyte subtype from perinatal clusters, as inferred from the consensus heatmaps. **e,** Dendrogram summarizing the inferred maturation steps of all astrocyte subtypes from multipotent or fate-restricted perinatal progenitor clusters. This highlights both convergence and divergence of cell fates, reflecting the complex scenario from which cerebellar astrocyte diversity arises postnatally. Note that the length of the connecting lines and the spatial arrangement of clusters in the dendrogram do not reflect the timing or duration of maturation steps, but were manually adjusted for clarity and to accommodate all elements within the figure layout. **f,** Expression levels of the top 10 DEGs of the five clusters predicted to be at the origin of the maturation trajectories. In situ hybridization data from the Allen Brain Atlas and immunofluorescence analyses suggest distinct spatial localizations of these clusters within the postnatal BG layer, cerebellar nuclei, or prospective white matter. Scale bars: 200 µm.

This strategy predicted that all BG subtypes originate from cluster 0 through maturing precursors in cluster 9, that WMA_*Npy* derive from cluster 17 via cluster 19 immature cells and that Vesic emerge from cluster 7 through their predecessors in cluster 20 (Fig. 3B). WMA_*Pgm5*, GLA_*arch* and CNA_*Gpc5* were all traced back to cluster 14 (Fig. 3B), suggesting that this cluster contains multipotent progenitors capable of giving rise to multiple astrocyte types. However, further analysis showed that cluster 14 comprises 3 cell subsets already displaying divergent commitments, each with distinct probabilities of differentiating into one of the three terminal subtypes (Extended Data Fig. 5G-I). An additional analysis to infer the origin of these 3 subsets (Extended Data Fig.5J; Fig. 3C) uncovered distinct progenitor sources: GLA_*arch* and WMA_*Pgm5* were predicted to derive from both cluster 1 and 4, while CNA_*Gpc5* originated exclusively from cluster 1. Hence, cluster 1 represents the only source of CNA_*Gpc5* through cluster 14, whereas the other two subtypes can emerge from either cluster 1 or 4. Applying a similar analysis to cluster 9 (Extended Data Fig. 5K), along trajectories of multiple BG subtypes, confirmed that all these subtypes originate with similar probabilities from cluster 0, highlighting homogeneous differentiation potential within these progenitor cells (Fig. 3C). Lastly, although hierarchical relationships among perinatal-enriched clusters could not be directly established (see Methods), clusters that ranked second-highest in initial analyses consistently rose to top positions in subsequent rounds (Fig. 3B, Extended Data Fig. 5D, dotted black lines), suggesting additional lineage transitions. In particular, cluster 4 may serve as a common progenitor for cluster 0—and thus for BG subtypes—as well as for GLA_*arch* and WMA_*Pgm5*.

To sum up, this analysis points to distinct perinatal progenitor clusters, each with unique differentiation potentials. Cluster 4 appears to contain multipotent progenitors giving rise to all BG subtypes, GLA_*arch* and WMA_*Pgm5*. By contrast, cells in cluster 0 are committed to the BG fate. Cluster 1 is the only source for CNA_*Gpc5*, though it can also differentiate into GLA_*arch* and WMA_*Pgm5*, but not BG. Clusters 17 and 7 exhibit more segregated trajectories, giving rise exclusively to WMA_*Npy* and Vesic, respectively. Of note, DEG analysis across these clusters and spatial mapping^37^ revealed diverse transcriptomic profiles and suggested separate spatial allocations in the perinatal cerebellum, well matching the inferred maturation trajectories (Fig. 3F, Extended Data Fig. 6E and Supplementary Table 6). Cluster 0 (BG source) expressed high levels of *Gria1* and *Gdf10* which are enriched in expanding BG at P4^23^ (Fig. 3F). Cluster 1 (CNA source) expressed *Slc6a11*, *Olig1* and *Olig2*, both highly expressed in the CN territory (Fig. 3F, Extended Data Fig 6F). Cluster 4 (multipotent progenitors of BG, GLA_*arch* and WMA_*Pgm5*) displayed elevated *Angpt1* and *Aqp4*, both enriched in the prospective WM (PWM; Fig. 3F); *Apoe*, shared by clusters 1 and 4, was likewise expressed in both CN and PWM (Extended Data Fig. 6B). Notably, the inferred PWM localization of cluster 4 multipotent progenitors aligns with the previously reported location of BG+GLA+WMA progenitors at birth^15^, confirming this cluster identity. Moreover, cluster 1, which showed elevated *Atp1b2*, mirrored the molecular profile of ACSA-2^+^/GLAST^+^ progenitors known to produce GLA and WMA but not BG^16^, consistent with cluster 1 inferred differentiation potential (Extended Data Fig. 6C,D). Interestingly, cluster 7, predicted to give rise to “glutamatergic-like” Vesic, showed high *Mytl1*^38^ and *Zmat4*, whose expression patterns match those of emerging glutamatergic cells (Extended Data Fig. 6B), suggesting a shared origin with excitatory neurons, distinct from other VZ-derived astrocyte subtypes. Due to limited data on DEG expression patterns, the spatial distribution of cluster 17 remained undetermined.

Finally, despite our integrative strategy did not identify any perinatal progenitor cluster common to both interneuron and WM astrocyte lineages^14^, likely due to low yield of astrocytes from this source, we hypothesized that cluster 11, with specific *Ptf1a* expression (Extended Data Figs. 5E, 6G) could contain such bipotent progenitors. This hypothesis was confirmed by genetic fate mapping using Ptf1a^CreERTM^ mice^39^ (Extended Data Figs. 5E, 6G) showing the generation of both ML interneurons and a smaller proportion of WM astrocytes.

Taken together, these results define the profile of CNA perinatal progenitors and reveal a complex mechanism underlying the emergence of astrocyte heterogeneity postnatally, driven by divergence and convergence of cell fates as well as by early fate-restricted paths (Fig. 3E).

### Cerebellar astrocyte heterogeneity unfolds from multiple lineages and shares embryonic origins with distinct other cell types

Having defined the maturation trajectories and fate potentials of perinatal astrocyte progenitors, we next sought to explore their embryonic origins and lineage relationships with other cerebellar cell types. To this end, we leveraged the clonal information contained in the P4-P5 dataset (Extended Data Fig. 3A), resulting from the injection at embryonic day 11 (E11) of a lipid nanoparticle (LNP)-encapsulated TrackerSeq library^40^. This transposon-based barcoding approach enables the high-throughput tagging of single embryonic progenitors, allowing for the retrospective identification of sister cells through scRNA-seq. Cells were grouped according to their assignment to distinct perinatal astrocyte progenitor clusters (i.e. “Slc1a3_cl*”) or to other cerebellar cell types, and lineage coupling scores were computed for all pairwise combinations (see Methods). As expected, hierarchical clustering of the pairwise correlations between coupling scores (Fig. 4A, Supplementary Table 7) revealed a lineage relationship between GABAergic interneurons and astrocyte progenitor clusters 0, 4, 17, and 19 (Fig. 4A-C), supporting a ventricular origin for BG, WMA_*Npy* and part of GLA_*arch* and WMA_*Pgm5*^14,15^. This was in line with spatial mapping onto an E16.5 ST dataset^41^, which localized clusters 0, 4, and 17 to the VZ and its adjacent cerebellar parenchyma (Fig. 4H). Intriguingly, these astrocyte progenitor clusters showed low coupling correlations with both clusters 1 and 7, suggesting a distinct developmental origin for CNA_*Gpc5* and Vesic.

**Fig. 4.**
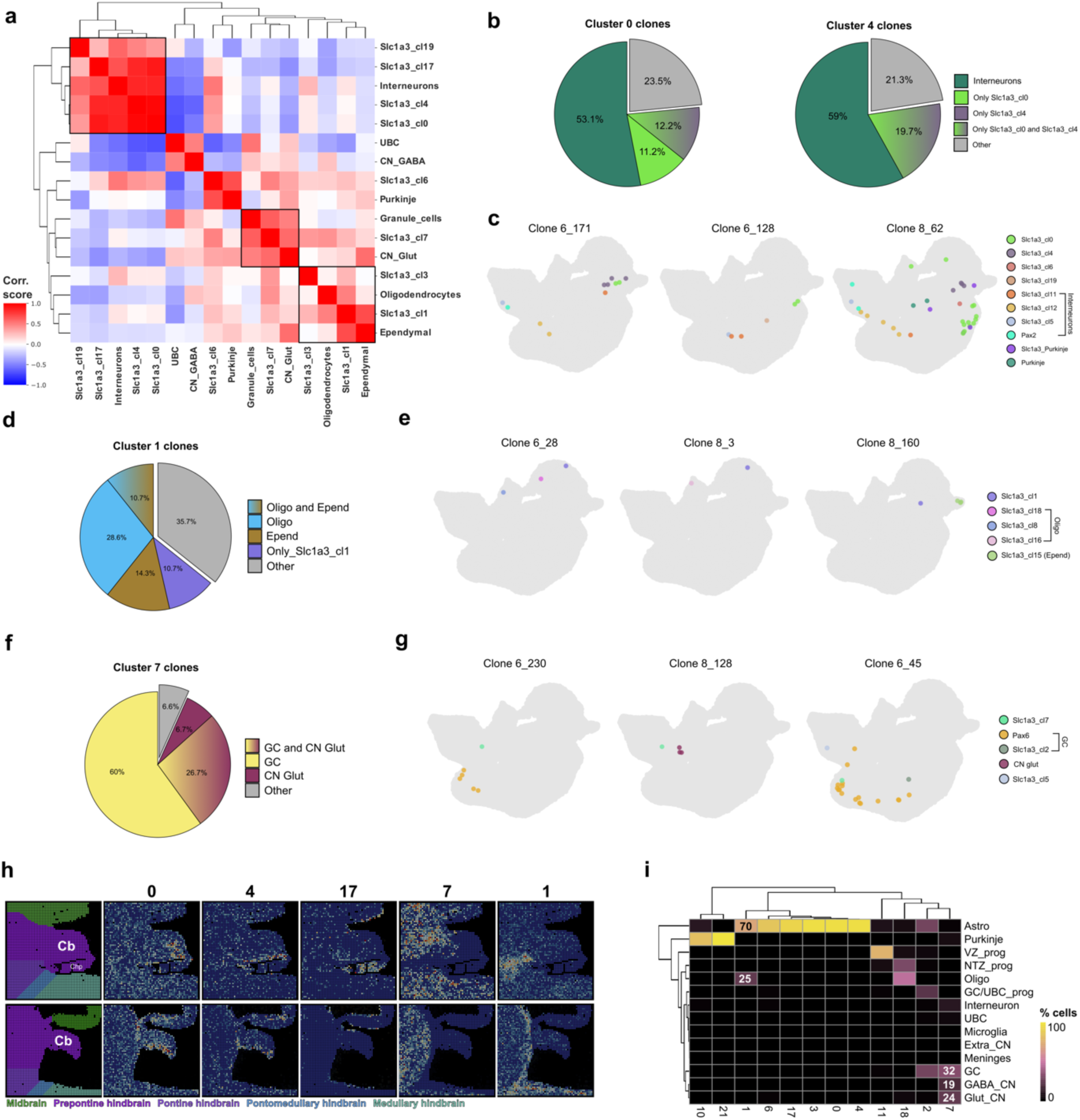
Lineage relationships and embryonic mapping of perinatal astrocyte progenitors. a, Heatmap of lineage coupling scores between pairs of P4 cells belonging to the perinatal astrocyte progenitor clusters (prefix “Slc1a3_”) or to other cerebellar cell lineages. Values range from 1 (red, highly coupled) to-1 (blue, anti-coupled). **b,** The pie charts show the composition of clones containing cells from cluster 0 (left) or cluster 4 (right). In both cases, more than 50% of the clones also contain interneurons. Notably, while 11% of cluster 0-containing clones are composed exclusively of cluster 0 cells, cluster 4-containing clones are always shared with other lineages, and 19% of them are shared with cluster 0, further supporting the multipotent nature of cluster 4 progenitors. **c,** Representative examples of clones shared between astrocyte progenitor clusters 0, 4 and 19 and cells along the interneuron lineage. Clone 8_62 (right) also contains cells in the Purkinje cell lineage (“Slc1a3_Purkinje” comprises cells in both “Slc1a3_cl10” and “Slc1a3_cl21” of the *Slc1a3* subset). **d,** The pie chart illustrates the composition of clones containing cells from cluster 1, highlighting the predominance of mixed clones with oligodendrocytes and/or ependymal cells. **e,** Examples of clones shared between cluster 1 and cells along the oligodendrocyte (left and middle plots) or ependymal (right plot) lineages. **f,** Pie chart showing the composition of clones containing cells from cluster 7, the vast majority of which also include granule cells (GCs, ∼87%) or glutamatergic CN neurons (∼33%). **g,** Representative examples of clones shared between cluster 7 and cells along the GC (left and middle plots) or glutamatergic CN neuron (right plot) lineages. **h,** Spatial mapping of perinatal astrocyte progenitor clusters onto two sections (hemisphere, top; vermis, bottom) from a E16.5 Stereo-seq dataset^41^. The first column shows the manual broad region annotation based on the Allen Brain Atlas, with corresponding color coding. The cerebellum (Cb) and choroid plexi (ChP) are labeled for visual reference. **i,** Heatmap showing the percentage of cells from each perinatal cluster of the *Slc1a3* subset (columns) that co-cluster with each of the distinct cerebellar cell type lineages in the integrated scRNA-seq dataset of the embryonic/early postnatal cerebellum. Relevant percentages are indicated within the corresponding squares.

**Figure 5.**
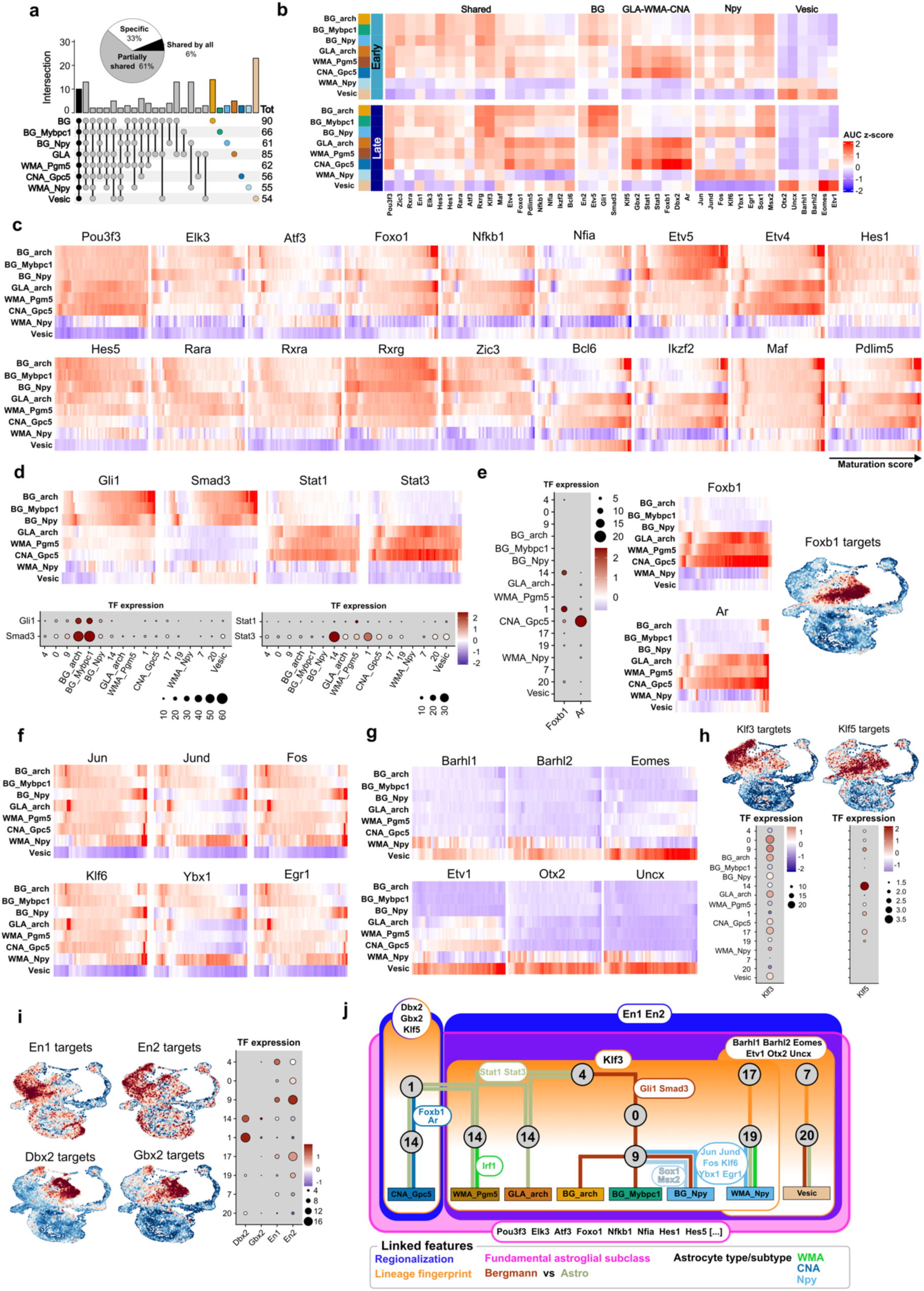
T**h**e **TR code underlying the emergence of cerebellar astrocyte heterogeneity. a,** UpSet plot illustrating the overlap of regulons identified by SCENIC across different astrocyte subtypes. The pie chart summarizes the proportion of regulons that are subtype-specific, partially shared among subtypes, or universally shared across all astrocyte subtypes. Totals on the right indicate the total number of regulons detected per subtype. **b,** Heatmap summarizing the activity of selected TRs (SCENIC-derived AUC z-scores) across the distinct trajectories in early (top) or late (bottom) maturation stages. TRs are grouped based on their activation pattern: shared across most trajectories, type-specific (BG, GLA-WMA-CNA) or subtype-specific (Npy, Vesic). **c,** Heatmaps showing the activity of TRs shared through the distinct trajectories. **d,** Selected TRs specific for Bergmann vs Astro. Heatmaps on the top show the activity of *Gli1* and *Smad3* (Bergmann-specific; left) or *Stat1* and *Stat3* (Astro-specific; right). Dot plots present the corresponding TR gene expression levels, consistent with their SCENIC-predicted activity. **e,** Selected TRs specific for CNA. Both gene expression (left) and predicted activity (middle) of *Foxb1* and *Ar* highlight specificity for CNA. UMAP plot shows the gene expression score of *Foxb1* regulon targets, enriched specifically in CNA progenitors within cluster 1. **f,** Heatmap showing the activity of TRs increasing through BG_*Npy* and WMA_*Npy* maturation trajectories. Note that the same TRs are active at the onset of the other trajectories and then gradually decrease. **g,** Heatmap showing the activity of TRs specific for the Vesic subtype(s). **h,** Target gene expression scores (top) and TR expression (bottom) of *Klf3* and *Klf5* showing complementarity between lineages stemming from cluster 4 or 1, respectively. **i,** Target gene expression scores (left and middle) and TR expression (right) of *En1/En2* or *Dbx2/Gbx2* showing complementarity between lineages stemming from cluster 4 or 1, respectively. **j,** Proposed model of the TR code orchestrating the emergence of cerebellar astrocyte types and subtypes.

On the one hand, cluster 1 appeared related to both ependymal cells and oligodendrocytes in the clonal data (Fig. 4A, D, E) and when projected onto an integrated sc/snRNA-seq dataset of the embryonic and perinatal cerebellum (Extended Data Fig. 7A-C), it was clustered with both astrocyte and oligodendrocyte lineages (70% and 25% of its cells, respectively; Fig. 4I). Moreover, its predicted localization at E16.5 (Fig. 4H) and the embryonic expression of GAT3 (*Slc6a11*, Fig. 3F; Extended Data Fig. 7D) were higher in the hindbrain than within the cerebellum, consistent with the extracerebellar origin of oligodendrocytes^13^. Supporting this developmental relationship, not only cluster 1 retained the expression of the oligodendrocyte-typical transcription factors *Olig1* and *Olig2* (Fig. 3F), but adult CNA_*Gpc5* also displayed a transcriptional profile closely resembling that of spinal cord and cortical astrocytes derived from *Olig2*-expressing progenitors^42,43^ (Extended Data Fig.7E-F), a well-established source of OPCs in the spinal cord^44^. Therefore, several lines of evidence pointed to a relationship between cluster 1 and the oligodendrocyte lineage.

On the other hand, cluster 7, the predicted source of Vesic, showed a strong lineage relationship with granule cells (GC) and with glutamatergic CN neurons (Fig. 4A, F, G; Extended Data Fig. 7G). In the integrated embryonic sc/snRNA-seq dataset, it clustered together with these excitatory populations (Fig. 4I) and it was predicted to localize within the nuclear transitory zone of the embryonic cerebellum (Fig. 4H), a transient developmental region where CN glutamatergic neurons differentiate^45^. Altogether, these findings further support a shared developmental origin between Vesic astrocytes and excitatory neurons, as previously discussed.

Altogether, these findings reveal diverse embryonic origins of cerebellar astrocyte subtypes. On the one hand, they confirm the VZ as the source of several astrocyte populations, extending to embryonic stages their lineage relationship with GABA interneurons, previously demonstrated only postnatally. On the other hand, they uncover novel associations with oligodendrocytes and excitatory neurons, suggesting that a subset of astrocytes arises from distinct germinative niches beyond the traditionally recognized VZ.

### The transcriptional logic driving the emergence of cerebellar astrocyte types and subtypes

To dissect the molecular mechanisms underlying cerebellar astrocyte maturation, we employed SCENIC^46^, which identified numerous transcriptional regulators (TRs) putatively active along multiple or specific trajectories (Fig. 5a, Supplementary Table 8). Manual curation and selection of TRs with predicted activity (see Methods) consistent with their expression pattern revealed a precise code orchestrating the emergence of cerebellar astrocyte heterogeneity at multiple levels (Fig. 5b, 5j, Supplementary Table 9).

A core set of regulators were shared through the distinct trajectories, either in the early or late maturation phases or through the whole process, indicating common molecular programs underlying the acquisition of features fundamental to the astroglial subclass (Fig. 5b,c). Consistently, many (*Pou3f3, Elk3, Atf3, Foxo1, Nfkb1* and *Nfia*) had documented roles in astrocyte differentiation, maturation and function^47–50^ or converged onto known developmental signaling pathways. The FGF-ERK-ETV axis (*Etv4*, *Etv5*), previously implicated in BG specification from radial glia^51^ was active also in other astrocyte types, with differential deployment of *Etv5* or *Etv4* in Bergmann vs Astro, respectively (Fig. 5b,c), pointing to a more general, though type-biased, role in cerebellar astrocyte induction and maintenance. Similarly, Notch signaling components (*Hes1*, *Hes5*) were prominently active at early stages of all trajectories except for Vesic, consistent with their established roles in gliogenic competence and BG monolayer formation^52,53^. Retinoic acid signaling (*Rara*, *Rxra*, *Rxrg*) also emerged as a shared regulatory network^54^. However, *Rara* and *Rxra* were more prominent at later stages of GLA_*arch* and WMA_*Pgm5* maturation, implying selective activation of this pathway in separate trajectories.

Notably, our analysis also revealed several new TRs, shared across trajectories (Fig. 5c, Extended Data Fig. 8b), that lacked prior association with astrocyte development. Specifically, *Zic3* was enriched at early stages, while *Bcl6*, *Ikzf2*, *Maf* and *Pdlim5* were found in more mature astrocytes. Their regulated genes are associated with critical astrocyte functions, such as modulation of glutamatergic synaptic transmission, positive regulation of neuron projection development, ion transport, and the regulation of blood-brain barrier permeability (Extended Data Fig. 8c). Hence, these are novel candidates involved in the functional maturation of cerebellar astrocytes.

We also observed trajectory-specific TRs, whose combination defines a transcriptional code guiding the differentiation of Bergmann vs Astro and, in turn, of distinct types and subtypes. Along all BG trajectories (Fig. 5d), we predicted a progressive increase in *Gli1*, consistent with its established role as a downstream effector of SHH signaling essential for BG maturation and maintenance of their molecular identity^24,55^. *Smad3*, similarly specific, may act downstream of *Zeb2* (or SIP1 - Smad-interacting protein 1-) in mediating TGF-β signaling during BG differentiation^56^.

Conversely, along the WMA_*Pgm5*, GLA_*arch*, and CNA_*Gpc5* Astro trajectories, high *Stat1* and *Stat3* implicate engagement of the JAK-STAT signaling pathway, consistent with its well-known role in cytokine-induced astrocyte differentiation^57^ (Fig. 5d). Notably, *Irf1* (Interferon regulatory factor 1), promoter of STAT1 DNA binding^58^, was high in WMA_*Pgm5* and WMA_*Npy* trajectories, pointing to a role in the acquisition of WMA type identity (Extended Data Fig. 8d). *Ar* was instead specific for the CNA_*Gpc5* trajectory (Fig. 5e), pointing to a role for androgen signaling in CNA maturation and/or maintenance. GO analysis of Ar targets revealed significant enrichment of genes involved in neurotransmitter uptake (Extended Data Fig. 8e), pointing to its contribution to the specialized homeostatic role of CNA within the cerebellar nuclei. Altogether, these TRs appear to guide cell fate decisions of cluster 4 and 1 multipotent progenitors.

Consistent with their role in shaping astrocyte type identity, many of the TRs described above were also active in the hybrid *Npy*-expressing and Vesic subtypes. *Etv5*, *Gli1* and *Smad3* were active in BG_*Npy*, while *Stat1*, *Stat3*, *Irf1* in WMA_*Npy* (Fig. 5c,d and Extended Data Fig. 8d), in line with their respective identities. Along the Vesic trajectory, shared regulators (*Foxo1*, *Nfia*, *Etv4*, *Etv5*, *Rara*, *Bcl6*, *Ikzf2*, *Maf* and *Pdlim5*; Fig. 5c), as well as Bergmann-(*Gli1*, *Smad3*) or Astro-specific TRs (*Stat1*, *Stat3*; Fig 5d) were active at late phases of maturation, when the fraction of mature astrocytes increases (Extended Data Fig. 8a). This combination further supports the hybrid identity of Vesic, comprising BG_*Kcnip4* and GLA_WMA_*Pde10a* (Fig 2f,g).

Other regulators were instead subtype-specific. Both BG_*Npy* and WMA_*Npy* displayed a highly specific predicted activity of several TRs associated with progenitor-state maintenance, including members of the JUN (*Jun, Jund*) and FOS (*Fos*) families, as well as *Klf6*, *Hey1*, *Ybx1*, *Atf4*, and *Egr1*^22^. These regulators, active in the early maturation phases of the other trajectories, increase later only in these two subtypes (Fig. 5f), consistent with their progenitor-like features. They also show elevated expression and activity of *Acaa1a*, a key enzyme in lipid catabolism, suggesting a reliance on alternative metabolic substrates to support their energetic demands (Extended Data Fig. 8f). Notably, BG_*Npy* also exhibited both strong expression and activity of *Sox1* and *Msx2* (Extended Data Fig. 8g). While *Sox1* is a known marker of specialized radial astrocytes in the hippocampus with demonstrated neurogenic and gliogenic potential in vivo^59^, *Msx2* shows enriched expression in human BG, based on snRNA-seq data^60^.

Taken together, SCENIC analysis identified combinations of transcriptional regulators specific for distinct astrocyte types and subtypes and therefore potentially orchestrating divergent, convergent or restricted fate decisions of cerebellar astroglial progenitors.

### Lineage-specific TRs contribute defining final astrocyte identity

We previously identified separate lineages (Fig. 4) contributing to astrocyte heterogeneity. Intriguingly, we found molecular traces of each embryonic origin persisting in the distinct trajectories.

Along the Vesic trajectory, several TRs characteristic of the cerebellar rhombic lip (RL) and its derivatives were highly active - including *Barhl1*, *Barhl2*, *Eomes*, *Etv1*, *Otx2*, and *Uncx* - (Fig. 5g). This regulatory signature supports the notion that Vesic represent a separate astrocyte lineage originating from the RL.

Along the trajectories originating from cluster 4 or cluster 1, which also correspond to distinct lineages (Fig. 3d), the two Krüppel-like factors (KLFs) *Klf3* and *Klf5* displayed striking complementary activity patterns (Fig. 5h). While other KLFs are known to be involved in cerebellar development and function^61,62^, the precise role of *Klf3* and *Klf5* remains to be elucidated. A similar pattern was followed by *En1* and *En2* vs *Dbx2* and *Gbx2* (Fig. 5i). On the one hand, *En1* and *En2* - homeobox genes master regulators in cerebellar development and known to persist up to late stages in many cerebellar cell types (including both VZ and RL-derivatives^63^) - were both expressed and predicted to be active along the trajectories originating from clusters 0, 4, 17 and 7, covering both VZ and RL progenies. On the other hand, *Dbx2* and *Gbx2* were high in cluster 1 and its derivatives. While *Gbx2* is known to be expressed in rhombomere 1 and required for cerebellar development during early embryogenesis but not at later stages^64^, both these genes are typically known for their role in hindbrain patterning^65,66^, consistent with the putative extra-cerebellar origin, in common with oligodendrocytes, of this cluster. A further link between cluster 1 and oligodendrocytes was represented by the activity of *Foxb1* (Fig. 5e), known to mark bipotent progenitors capable of generating both oligodendrocytes and astrocytes in the medulla and thalamus^67^.

In parallel, *Egfr*, implicated in glial lineage plasticity and in the regulation of astrocyte-to-oligodendrocyte conversion and vice versa^68^, was expressed in both cluster 1 and 18, at the onset of the oligodendrocyte lineage. The *Fli1* regulon also showed a similar pattern of activity in both clusters (Extended Data Fig. 8h). Given its previously uncharacterized role in glial development, *Fli1* may represent a relevant target to explore the potential switch between astrocyte and oligodendrocyte fates from a common progenitor.

Overall, SCENIC analysis allowed to uncover a TR code underlying the emergence of cerebellar astrocyte heterogeneity (Fig. 5j), which comprises long lasting traces developmentally inherited from their progenitor sources and that contribute to defining their final identity.

## Discussion

This study provides a comprehensive definition of molecularly distinct cerebellar astrocyte types and subtypes, and reconstructs *in silico* the developmental mechanisms underlying their emergence, revealing multiple maturation trajectories. These trajectories encompass fate divergence, convergence and restriction, involving multiple progenitor sources that share developmental origin with other cerebellar cell subclasses. Additionally, this study infers the TR modules delineating the unfolding of astrocyte lineages, up to the acquisition of type and subtype identities. Overall, these findings indicate that while environmental cues predominantly guide the acquisition of terminal astrocyte molecular fates, lineage specific factors constrain progenitor potency and impart defined traits in astrocyte subtypes.

Sc/snRNAseq and ST identified established and novel adult astrocyte profiles, including four BG (BG_*arch*, BG_*Mybpc1*, BG_*Npy* and BG_*Kcnip4*), two GLA (GLA_*arch* and GLA_*Pde10a*) and three WMA subtypes (WMA_*Pgm5*, WMA_*Npy* and WMA_*Pde10a*), along with new molecular markers for their discrimination. Notably, BG_*arch* and BG_*Mybpc1* displayed complementary distributions across anterior and posterior lobules, respectively, consistent with prior observations^19,32^. This spatial segregation may depend on involvement in distinct circuitries driven by different afferents, potentially leading to molecular divergence^17^. Further on functional compartmentalization, *Npy* enriched BG subtype (BG_*Npy*) could correspond to the striped *Npy-Gfp*-labeled BG^69^. However, investigation of ST datasets (not shown) found no evidence of parasagittal BG banding. Thus, although current molecular profiling effectively captures anteroposterior BG heterogeneity reflective of distinct circuitries, it does not identify transcriptional substrates corresponding to biochemical and functional parasagittal neuron topography^70^. Yet, BG_*Npy* exhibited distinctive characteristics, and shared with WMA_*Npy* traits reminiscent of quiescent neural stem-like cells. These shared features partly recapitulate those of a subset of astrocytes in the neocortical WM/corpus callosum, known to actively support astrogenesis^22^. Moreover, BG_*Kcnip4* and GLA_WMA_*Pde10a* exhibited a unique molecular profile denoting glutamatergic gliotransmission. Their true astrocyte identity is supported by expression of canonical Bergmann or Astro markers, activation of TR involved in astrocyte differentiation, and consistent detection across diverse datasets (adult and postnatal snRNA-seq; ST dataset completely devoid of non-astrocytic cells). Of note, these clusters closely resemble glutamatergic astrocytes found in the hippocampus and substantia nigra^29^, aligning with formerly reported gliotransmission in the cerebellum^71^.

GO analysis captured a broad functional divergence between the two Bergmann and Astro groups. While Bergmann main functions in maintaining cerebellar structural integrity, regulating glutamatergic inputs, and relying on Shh for their cellular identity^13^ were confirmed, Astro roles, much less characterized, were found primarily related to energy homeostasis and neuroprotection. Yet, since BG are known to display high glucose transport and metabolic activity^72^, the enrichment of energetic pathways in Astro may reflect compensatory transcriptional upregulation in response to a lower cellular density compared to BG, rather than indicating an exclusive specialization. In contrast, the stronger neuroprotective signature in Astro could point to a genuine functional enrichment. Oxidative stress and lipid peroxidation are more pronounced in the PCL and ML across multiple disease models^73–75^ and amyloid plaques in Alzheimer’s disease are predominantly found in the ML^76^. This suggests a higher resilience of other cerebellar regions possibly supported by enhanced astrocyte properties.

The *in silico* reconstruction of the maturation trajectories revealed a surprisingly complex picture of astrocyte generation from multiple perinatal sources with divergent, convergent, or restricted trajectories. Along these trajectories, TRs appeared to operate at different stages according to combinatorial modules coding regional patterning and regulating lineage-specific, subclass-specific and astrocyte type/subtype-specific features (Fig. 5j, Supplementary Table 9). First components of the TR code were the regional patterning factors *En1/En2* or *Dbx2*/*Gbx2.* These factors are active in VZ-and RL-derived cerebellar lineages or within the hindbrain at the embryonic stages when astrogliogenesis is initiated and were associated with distinct trajectories in our data. Further TR modules progressively overlap, delineating VZ vs RL origins and contributing to the acquisition of specific astrocytic traits.

Fate divergence towards BG subtypes, GLA_*arch* and WMA_*Pgm5* emerged from multipotent cells (cluster 4), well matching formerly identified PWM progenitors^15^, now defined by the expression of *Angpt1* and *Aqp4*. Clonal data support that these lineages share common embryonic VZ progenitors with GABAergic interneurons, thereby extending former evidence for postnatal bipotent progenitors^14^ and confirming former genetic studies^77,78^. All of the aforementioned divergent trajectories are delineated by the expression and activity of the cerebellar patterning factors *En1* and *En2*, consistent with their origin from the cerebellar VZ^15^, and of *Klf3*, whose activity pattern, complementary to that of *Klf5* (active in *Dbx2/Gbx2-* associated derivatives), warrants further investigations. Along time, these factors are associated with fundamental astroglial fate determinants (e.g. *Hes1, Hes5, Nfia, Foxo1*), and specific promoters of either Bergmann or Astro fates. Bergmann trajectories exhibited active Shh and TGF-β signaling whereas Astro displayed cytokine responses (Supplementary Table 9). In addition, Irf1 association with WMA fates suggests a role for interferon in the maturation of this type.

Notably, as mentioned before, the BG_*Npy* trajectory was distinctively enriched for TR particularly active in progenitors (transcription factors of the AP-1 family *Jun*, *Jund*, *Fos*, *Atf4*), suggesting that this subset may possess particularly plastic properties^22^. Similar features were shared by WMA_*Npy* which nevertheless showed an independent restricted origin from cluster 17. While clonal tracing supported a common ancestry with other VZ-derived astrocytes (i.e., cluster 4 derivatives), at the onset of the WMA_*Npy* trajectory several RL-typical transcription factors such as *Barhl1*, *Barhl2*, *Eomes*, *Etv1*, *Otx2* and *Uncx* were found activated. This may suggest a specific ontogenic origin, possibly within the “posterior transitory zone”, a VZ region immediately anterior to the morphologically defined RL where Atoh1 lineage progenitors possess an astrogenic potential^79,80^.

Another restricted trajectory moved from cluster 7 to form glutamatergic Vesic astroglia, comprising the BG_*Kcnip4* and GLA_WMA_*Pde10a* subtypes. Sustained expression and activity of the RL-typical transcription factors mentioned above, combined with *En1/En2*, together with clonal analysis and spatial mapping all point to their RL origin. Interestingly, astroglia fate induction appears to occur through the rather late activation and expression of general astrocyte or type-specific TRs. The contribution of the RL to astrogenesis has long been debated, with conflicting evidence both supporting and opposing this possibility^81^. Our data now provide compelling support in favor of a RL contribution. This offers a novel perspective on lineage relationships and raises the possibility that similar ontogenetic programs may give rise to analogous astrocyte populations in other brain regions^29^.

Further, additional multipotent progenitors were identified in cluster 1. However, within this developmental trajectory, both the fate potency and inferred TRs appeared to diverge from those characterizing cluster 4. Cluster 1 was predicted as unique source of CNA, in parallel with divergence onto GLA_*arch* and WMA_*Pgm5,* also converging from cluster 4. Whether this reflects full phenotypic convergence, as described for specific neuronal types^82^, remains unclear. One lineage may eventually dominate, or lineage-specific traits may persist in ways not captured by transcriptomic profiling. Supporting the latter hypothesis, GAT-3 immunostaining revealed a few positive GLA and WMA in the deep cerebellum, in addition to its expression in CNA (Extended Data Fig. 6f). Since *Slc6a11* (GAT-3 gene) is specifically enriched in cluster 1, this suggests that at least a small subset of GLA and WMA can be traced back to this origin. Notably, and in contrast to the *En1/En2*-defined trajectories, cluster 1 and its derivatives exhibited predicted activity of the hindbrain patterning factors *Dbx2* and *Gbx2*, in association with *Klf5* as well as general astroglial and Astro-specific TR, consistent with its fate potency. Ultimately, additional regulation by *Foxb1* and *Ar* seems to operate towards CNA.

Further analyses of cluster 1 also suggested formerly unknown origins for CNA and the other diverging trajectories. Spatial mapping onto embryonic ST datasets, expression of hindbrain TRs, its lineage relationship with oligodendrocytes, as well as CNA similarity with brainstem astrocytes^25^ all argue against a VZ origin. Indeed, most cerebellar oligodendrocytes originate from an extra-cerebellar source^77,83,84^ and CNA were rarely labeled in VZ-targeted *in utero* electroporation clonal analyses^15^. Moreover, this supports earlier suggestions of bipotent astrocyte-oligodendrocyte progenitors in the cerebellum, based on indirect *in vivo* and *in vitro* evidence of fate shifts between these glial types upon Bmi1 deletion or BMP pathway inhibition^85^. Molecular traces of such bipotency can also be found in the CNA trajectory.

Notably, most of the TR predicted to regulate astrocyte type identity are downstream of diffusible extrinsic signals (e.g., SHH, TGF-β, IFN, cytokines, androgen, retinoic acid), highlighting the pivotal role of the local environment in shaping astrocyte identity. This suggests that spatial and/or temporal gradients of these signals across cerebellar layers may guide fate acquisition and maintenance, as shown for SHH^24^. Consistently, several of these signals exhibit dynamic secretion patterns during cerebellar development^86–89^ and astrocyte types follow distinct temporal differentiation patterns^15^. In contrast, TR downstream of contact-dependent pathways (e.g. Notch) were less prominent, possibly due to the transient or spatially restricted nature of their activation, which may limit their detection by SCENIC.

On the other side, lineage-related factors act cell-autonomously to restrict the fate potency, as exemplified by transplantation studies: when grafted into the developing cerebellar cortex, both cortical astrocyte progenitors and ACSA2+ cerebellar astrocyte progenitors (i.e. cluster 1) fail to generate BG, retaining their original fate potency^16,90^. These findings underscore that extrinsic signals alone are insufficient to impose astrocyte identities in the absence of a pre-existing intrinsic competence, like the cerebellum-specific patterning regulators En1/En2.

By achieving an unprecedented resolution on cerebellar astrocyte heterogeneity and decoding the underlying regulatory landscape, this study provides a conceptual advance in understanding how astrocyte heterogeneity is developmentally achieved. These insights lay the foundation for exploring whether similar regulatory programs contribute to astrocyte specialization throughout the brain, with important implications for understanding their physiological functions and disease responses.

## Materials and Methods

### Experimental animals

C57BL/6J (Charles River Laboratories), C57BL/6JRj (Janvier), CD1 (Charles River Laboratories), hGFAPeGFP (FVB/N-Tg(HGFAP::EGFP^91^) and Ptf1a^CreERTM^ ^39^ crossed with R26R^EYFP/EYFP^ (R26R^YFP 92^were housed at two to five animals per cage under a 12 h–12 h light– dark cycle (lights on from 07:00 to 19:00) at a constant temperature (22 ± 2 °C) and humidity (55 ± 10%) with ad libitum access to food and water. Mice were used at 2-3 months of age and at different postnatal (P) times as indicated in the Results. Mice of either sex were included. The day of vaginal plug detection was defined as E0, and the day of birth was considered as P0.

All animal protocols in the present study were approved by the Swiss Federal and Cantonal authorities, by the Council Directive of the European Communities (2010/63/EU) or by the Government of Upper Bavaria (Germany) or by the Italian Ministry of Health.

### Tamoxifen injections in postnatal mice

To induce Cre recombination in Ptf1a^CreERTM^ × R26R^YFP^ P0 mouse pups, we administered tamoxifen (Tx; 20 mg/ml solution dissolved in corn oil) via subcutaneous injection (600 μg/pup).

### LNP-TrackerSeq library preparation

The TrackerSeq library, a piggyBac transposon-based system compatible with the 10x Genomics single-cell transcriptomic platform, was prepared following the protocol described in Bandler et al.^40^. This library enables *in vivo* lineage tracing by integrating synthetic nucleotide barcodes into the mouse genome. Briefly, the piggyBac donor plasmid (Addgene #40973) was modified to incorporate a Read2 partial primer sequence within the 3′ UTR of eGFP to facilitate retrieval in the 10x pipeline, as well as a sucrose selection system to eliminate empty plasmids. The lineage barcode oligo mix was cloned downstream of the Read2 partial primer sequence via Gibson Assembly. The assembled constructs were then transformed into E. coli, expanded, and purified using standard column-based methods. The resulting plasmid library was quality-checked by high-depth sequencing to assess barcode diversity and overrepresentation.

For *in vivo* delivery, the purified TrackerSeq library was incorporated into lipid nanoparticles (LNPs). LNPs were formulated to encapsulate three plasmids: pCAG-Trackerseq, pCAG-GFP-KASH, and pEF1a-PBase, thereby enabling robust and heritable integration into the progenitor genome. Plasmid DNA was combined to a total of 20 µg. LNP encapsulation was performed using an automated microfluidic system following the manufacturer’s recommended protocol, which combines the plasmid mixture with a specific buffer and a proprietary lipid nanoparticle mix conceived for progenitor targeting. The resulting LNP-encapsulated plasmids were stored at 4°C. For quality control, LNP size distribution was determined by Dynamic Light Scattering (DLS) using a ZetaSizer (Malvern Panalytical). Samples were diluted in PBS, and measurements were performed at 25°C. Optimal LNP preparations typically exhibited a single size peak between 100-120 nm.

### LNP-TrackerSeq injections in mouse embryos

Timed-pregnant mice were deeply anesthetized throughout the surgery duration with isoflurane/oxygen and a small abdominal incision was performed to access the uterine horns. Embryos were constantly maintained moist with physiological saline. On embryonic day (E) 11 or E14, 70nl of the LNP-TrackerSeq library were injected into the third ventricle of the embryos. 50ul of Temgesic was injected 30 minutes before the surgery and 6 hours after surgery for analgesia.

### Sample collection for sc/snRNA-seq

#### Nuclei collection from the adult deep cerebellum

Single nuclei from the adult deep cerebellum were obtained as follows. Brains from two P56 hGFAPeGFP or C57BL/6JRj mice were freshly dissected and immediately transferred to cold artificial cerebrospinal fluid (ACSF). Subsequently, they were embedded into 4% Agarose Low Melt at 37°C and cut parasagittally into 600-μm-thick sections. Sections where the CN were clearly visible were selected and the deep cerebellum, containing both the CN and the deep WM, was macrodissected with a microsurgical knife. Tissues from both animals were pooled together and further processed for nuclei isolation following the 10X Chromium recommended protocol CG000366 (Rev A) with small adaptations. Nuclei suspensions from hGFAPeGFP mice were incubated for 30min at 4°C with a PE-coniugated anti-NeuN antibody (Milli-Mark® anti-NeuN-PE, clone A60, Sigma, 1:500) to label neuronal nuclei and subsequently subjected to FACS sorting (BD FACSAria III Cell Sorter). For the FACS gating, we used side scatter versus forward scatter and 7-AAD staining (7-Aminoactinomycin D, ThermoFisher) to distinguish single nuclei from debris and doublets or aggregates. NeuN-PE unstained cells were therefore collected to enrich non-neuronal nuclei. After FACS, nuclei were suspended in FACS buffer at a final concentration of 3000 cells/µl and loaded into the 10x Chromium V3 system. Reverse transcription and library generation were performed according to the manufacturer’s protocol.

#### Cell collection from the adult deep cerebellar WM

Single cells of the adult deep cerebellar WM were isolated from 8-12 weeks-old C57BL/6J male mice (12 mice in total, in two independent experiments) as previously described^22^. In brief, parasagittal slices with 1mm intervals were obtained by placing the freshly dissected cerebella onto an acrylic adult mouse brain matrix slicer and the deep WM was manually dissected through examination under a stereomicroscope. This was achieved excluding the CN and the GL to avoid contamination from the surrounding GM territories. Single cells were isolated from the dissected tissues using the Papain Dissociation System (Worthington Biochemical Corporation, 60 minutes incubation) followed by the Dead Cell Removal kit (Miltenyi Biotec) and Myelin Removal Beads II (Miltenyi Biotec) according to the manufacturer instructions. Single cells were eluted and suspended in 1X PBS with 0.04% BSA at a final concentration of 1000 cells/µl for subsequent processing using the Single Cell 3’ Reagent Kits v3.1 (10x Genomics) following the manufacturer instructions.

#### Cells and Nuclei collection from the postnatal cerebellum

Single nuclei were isolated from postnatal cerebella ranging from P0 to P30. Brains from C57BL/6JRj, CD1 or hGFAPeGFP mice of both sexes (31 in total, in 22 independent experiments, Supplementary Table 1) were freshly dissected and immediately transferred to cold artificial cerebrospinal fluid (ACSF).

In 12 experiments, brains were directly processed for nuclei isolation following the 10X Chromium recommended protocol (CG000366 Rev A) with minor adaptations.

In the remaining 10 experiments, brains were embedded into 4% low-melting agarose at 37°C and cut parasagittally into 600-μm-thick sections. From each section, the anterior, central, and posterior lobules, as well as the deep cerebellum, were macro-dissected using a microsurgical knife. Adjacent sections containing the same anteroposterior (AP) regions were pooled together, with separate processing for those collected from the vermis and from three consecutive mediolateral (ML) portions of the two hemispheres (the same portions of the two hemispheres were pooled together). This approach ensured that spatial information along both AP and ML axes regarding the nuclei origin was preserved. All samples were subsequently processed for nuclei isolation as described above.

To enable multiplexing of samples from distinct regions, ages or biological replicates (see Supplementary Table 1), single nuclei were stained with TotalSeqA™ HashTag Oligonucleotide (HTO) antibodies (BioLegend) following the manufacturer’s protocol with minor adaptations. Briefly, nuclei were resuspended in staining buffer, incubated with a TotalSeqA™ antibody cocktail for 30 min at 4°C, and washed twice with PBS. The stained nuclei from distinct samples were merged into a single multiplexed sample, filtered through a 40 µm strainer, and loaded into the 10x Chromium V3 system.

Single cells were isolated from P4, P5 or P23 CD1 mice injected with the LNP TrackerSeq library at E11 or E14 (17 in total in 5 independent experiments, Supplementary Table 1). Briefly, cerebella were collected and maintained in Leibowitz medium supplemented with 5% FBS. Tissue dissociation was performed using the papain-based system (Worthington), following the manufacturer’s protocol. eGFP labeled cells were then isolated via flow cytometry using a BD FACSAria III Cell Sorter. Non-eGFP-expressing brain tissue served as a negative control to define background fluorescence in all FACS experiments. The sorted cells were processed using the Single Cell 3’ Reagent Kits v3.1 (10x Genomics) following the manufacturer instructions.

### Construction of sc/snRNA-seq, TotalSeqA™ HTO and TrackerSeq libraries

Sc/sn barcoding and library construction were performed using the Chromium Single Cell 3’ Reagent Kits (v3 chemistry) and the Chromium Controller instrument (10x Genomics), following the manufacturer’s instructions.

The amplification of the TotalSeqA^TM^ HTO library was performed according to manufacturer instructions. Briefly, the cDNA amplification reaction was modified by adding 0.2 µM of an HTO-specific additive primer to enhance the yield of HTO-derived cDNA products. The reaction mix consisted of 50 μl Amplification Master Mix, 7 μl nuclease-free water, 5 μl cDNA additive, 2 μl cDNA primer mix, and 1 μl of the HTO-specific additive primer, for a total volume of 65 μl. Following cDNA amplification, HTO-derived cDNA (∼180 bp) was separated from mRNA-derived cDNA (>300 bp) using a two-step sparQ or SPRI bead-based selection (0.6X for separation, followed by 2X purification of the supernatant containing the HTO fraction). The purified HTO fraction was then amplified using KAPA HiFi HotStart ReadyMix (2X), with a total reaction volume of 100 μl containing 45 μl purified HTO cDNA, 50 μl KAPA HiFi HotStart ReadyMix, 2.5 μl Illumina TruSeq D70*_s i7 index primer (10 µM), and 2.5 μl SI-PCR oligo (10 µM). The amplified HTO library was further purified using a 1.6X sparQ or SPRI bead-based purification.

The amplification of the TrackerSeq barcode library from RNA was carried out as previously described^40^, using the Q5 Hot Start High-Fidelity 2X Master Mix (NEB, #M094S) according to the standard NEB protocol. Each 50-μl PCR reaction included 25 μl of Q5 High-Fidelity 2X Master Mix, 2.5 μl of a 10 μM P7-indexed reverse primer, 2.5 μl of a 10 μM i5-indexed forward primer, 10 μl of molecular-grade H₂O, and 10 μl of cDNA as the template.

### Sequencing and Data Processing

**I**llumina sequencing libraries were sequenced with HiSeq 4,000 or NovaSeq 6,000 after quality assessment with the Bioanalyzer (Agilent) with an average read depth of 20,000-30,000 raw reads per cell. Sequencing reads in FASTQ format were aligned to the mouse reference transcriptome (mm10-2.1.0) and converted into Unique Molecular Identifier (UMI) counts using Cell Ranger software with default settings (10x Genomics, version 6.0.1). Following sequencing, the applied HTO sequences were utilized for bioinformatic demultiplexing of nuclei and the resolution of individual samples.

For TrackerSeq, preprocessing of R2 FASTQ reads involved trimming sequences flanking the lineage barcodes (BC). Barcodes shorter than 37 base pairs were discarded. Cell barcodes (Cell) were extracted from the corresponding Seurat dataset to generate a whitelist, which was then used to incorporate cell barcodes and UMIs into the read names of the lineage barcode FASTQ files. These processed FASTQ files were further analyzed to construct a sparse matrix in CSV format, where rows represented individual cells identified by unique cell barcodes, and columns corresponded to lineage barcodes. Only Cell–UMI–BC triplets supported by at least 10 reads and Cell–BC pairs with a minimum of 6 UMIs were retained for further analysis. CloneID assignment to cell barcodes was performed using Jaccard similarity and average linkage clustering, following the approach described by Wagner et al.^40,93^.

### General data processing and quality control

Sc/snRNA-seq datasets were pre-processed with CellBender or SoupX to remove technical noise and ambient RNA, improving the overall accuracy and quality of the data^94,95^.

Subsequent processing was performed using the Seurat pipeline (versions 5.0.1 or 5.0.2). Raw count matrices were imported into R (versions 4.3.2 or 4.3.3) and converted into Seurat objects using the CreateSeuratObject() function. Cells were filtered based on dataset-specific thresholds for gene detection (minimum and maximum number of genes per cell) and mitochondrial read percentage.

### Publicly available sc/snRNA-seq datasets

Count matrices and associated metadata of adult mouse cerebellum sn/scRNA-seq datasets were downloaded from https://singlecell.broadinstitute.org/single_cell/study/SCP795 (Kozareva et al.^18^), https://www.ncbi.nlm.nih.gov/geo/query/acc.cgi?acc=GSE246717 (Yao et al.^20^), and https://www.ncbi.nlm.nih.gov/geo/query/acc.cgi?acc=GSE160471 (Kebschull et al.^21^). Embryonic and postnatal available datasets were downloaded from https://www.ebi.ac.uk/ena/browser/view/PRJEB23051 (Carter et al.^32^), https://www.ncbi.nlm.nih.gov/geo/query/acc.cgi?acc=GSE118068 (Vladoiu et al.^33^). The Sepp et al.^19^ dataset, used for both our adult and developmental studies, was downloaded from https://cellxgene.cziscience.com/collections/72d37bc9-76cc-442d-9131-da0e273862db

### Sc/snRNA-seq dataset integration

#### Adult sc/snRNA-seq integrated dataset

The distinct datasets were merged together in a single Seurat object, which was then split into individual layers, each corresponding to one dataset of origin (batch). Normalization using the SCTransform() function (regressing out known sources of unwanted variability, like sequencing depth and mitochondrial transcript proportion) and Principal Component Analysis (PCA) were performed on each layer. Then, to enable efficient integration of large-scale datasets, we applied the Sketch integration approach implemented in Seurat v5. From each dataset (layer) a representative subset of cells was selected using leverage score-based sampling. These sketched cells were integrated using a canonical correlation analysis (CCA)-based workflow to define a shared low-dimensional space. The full datasets were then projected onto this space, allowing harmonized analysis across all cells while minimizing memory usage. Following integration, clusters were identified using the FindNeighbors() and FindClusters() functions in Seurat. The resulting clusters were annotated based on the expression of literature-based cell type-specific marker genes as shown in Extended Data Fig. 1b.

To analyze adult astrocytes, an “astrocyte score” was calculated using the Seurat function AddModuleScore(), based on the combinatorial expression of the astrocyte-specific marker genes *Slc1a3, Slc1a2, Aldh1l1*, and *Aqp4*. Cells/nuclei with an astrocyte score greater than 1 were subsetted for further analysis. This subset was then subjected to the same computational workflow described above, including batch-wise integration using CCA in Seurat V5, to generate the final integrated dataset of adult cerebellar astrocytes. This dataset did not comprise any cells from Kebschull et al.^21^, since the authors sorted *NeuN+* neuronal nuclei, thereby excluding non-neuronal cell types, including astrocytes.

#### Postnatal sc/snRNAseq integrated dataset

For the postnatal dataset, we adopted a similar Seurat v5 integration strategy but without sketching the data. Instead, the full dataset was merged into a single Seurat object and split into layers corresponding to homogeneous experimental cohorts - defined by shared technical parameters such as 10X chemistry, mouse strain, and sample type (e.g., nuclei or cells) - to account for batch effects while preserving biological variation. Normalization was performed independently per layer, followed by CCA-based integration, and clustering.

*Slc1a3*-expressing cells were subsetted and re-integrated following the same workflow adopted for the whole dataset. The same approach was also used for the integration of our perinatal *Slc1a3*-expressing cells with the embryonic-perinatal cells of publicly available datasets^19,32,33^, each treated as separate experimental cohort.

### Spatial mapping onto ST datasets

To infer the spatial location of the clusters of either the adult or postnatal datasets, identity transfer of the cluster identities was performed on distinct ST datasets.

#### Visium

The sketched representative subset of the adult whole cerebellum dataset (see above), the adult astrocyte datasets or the postnatal *Slc1a3* subset were integrated with the adult mouse brain sagittal slice “posterior 1” of the Visium ST dataset “stxBrain” available on Seurat (Mouse Brain Serial Section 1 (Sagittal-Posterior), Spatial Gene Expression Dataset by Space Ranger 1.1.0 (2020, June 23)). An “anchor”-based integration workflow in Seurat was applied, which enables the probabilistic transfer of annotations from a reference to a query set^96^. The spatial reference dataset and the adult astrocyte dataset were normalized using the SCTransform function, which builds regularized negative binomial models of gene expression. Dimensionality reduction using the RunPCA function and label transfer using the functions FindTransferAnchors and TransferData were performed. This procedure outputs, for each spatial spot, a probabilistic classification for each of the sc/snRNAseq-derived cell states. These predictions were added as a new assay in the Seurat object for visualization using the function SpatialFeaturePlot.

#### Merfish

The adult astrocyte clusters were aligned with 16 sagittal cerebellar slices of the adult mouse cerebellum^20^. Prior to alignment, cells in each slice were subjected to cluster analysis with Seurat with FindClusters at resolution 0.5^96^. Cells in slices were then scored based on the combined expression of astrocytic markers (*Aqp4*,*Aldh1l1*,*Sox9*) with the Seurat function AddModuleScore. Clusters containing the majority of cells with high expression of those markers and cells with scores higher than 2 standard deviations from the mean of the astrocytic score were subsetted and then used for alignment with the adult astrocyte dataset. To align spatial cells with the adult astrocyte dataset an approach based on a loss function of cosine similarity between single cell and spatial data was performed with the Python package Tangram^97^. Clusters from the adult astrocyte dataset were fitted on each cerebellar slice with the tangram function map_cells_to_space based on the normalized expression of a set of training genes. Training genes were defined with the Scanpy^98^ function tl.rank_genes_groups and the tangram function pp.adatas, this process scored each cell in the slices based on their probability of belonging to the clusters of the adult astrocyte dataset. Cells in slices were assigned to a cluster identity when their score for belonging to the cluster was higher than 2 standard deviations from the mean of the identity score in the slice.

#### Stereo Seq

The subset of *Slc1a3*-expressing cells from postnatal dataset were aligned with two sagittal mouse brain sections from embryonic day (E) 16.5^41^ and one sagittal mouse brain section from postnatal day 7^37^. Prior to alignment cells belonging to the developing cerebellum and nearby structures were subset and then used for alignment. Astroglial-like progenitors and maturing astrocytes were aligned with the spatial slices with the package Tangram^97^ as described in the Merfish paragraph above.

### Layer enrichment and lobule enrichment analyses

#### Merfish

For the layer enrichment analysis, each cell in each Merfish cerebellar slice was assigned to one of four layers: cerebellar nuclei (CN), white matter (WM), granular layer (GL) and molecular layer/Purkinje cell layer (ML/PCL). This manual annotation was performed using a custom R script, based on both cell density patterns observed in the Merfish images and, when available, the expression of regionally enriched marker genes (*Sox10* for WM, *Calb1* for ML/PCL, *Slc17a7* for GL, while for cerebellar nuclei a pre-existing annotation was adopted). For the cerebellar nuclei, a pre-existing annotation was used. Cells located in extracerebellar regions or in the choroid plexus were excluded from the layer assignment and downstream analysis.

For the lobule enrichment analysis, a similar manual curation strategy was applied to assign cells in each MERFISH slice to the corresponding cerebellar lobule. In both analyses, enrichment scores for each cluster identity *i* within each layer or lobule *j* were computed as:

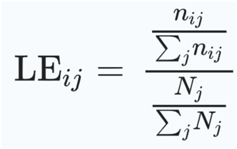

where n_ij_ is the number of cells from cluster *i* assigned to layer/lobule *j*, and *N_j_* is the total number of cells in layer/lobule *j* (across all cell types).

#### Kozareva et al. snRNA-seq dataset

To compute the lobule enrichment scores for single-nucleus transcriptomic profiles, astrocytes from the dataset by Kozareva and colleagues^18^ were subset from the integrated adult sn/scRNA-seq dataset. The original metadata indicating the lobular origin of each nucleus was combined with our cluster identity annotations. Lobule enrichment scores were then calculated using the same formula described above, where the number of nuclei belonging to each cluster and lobule was normalized by the total number of nuclei per lobule.

### Cell types/subtypes annotation

To assign cell type identities across the whole cerebellum datasets, we used a deep neural network classifier, implemented in PyTorch, following the strategy described by De Ceglia et al.^29^. Briefly, a multiclass prediction model was trained using pre-annotated single-cell transcriptomic data. Cell types with fewer than 50 annotated cells were excluded from the training set to ensure robust classification. The dataset was randomly split into training (80%) and testing (20%) subsets.

The network architecture consisted of four fully connected layers. Hardtanh was used as an activation function in the first layer, and ReLU for the subsequent layers. Dropout was applied during training (50% at the input of the first layer, 30% at the following layers) to prevent overfitting. Additionally, the weights of the first layer were normalized to unit norm for each node, and input normalization was adjusted accordingly. Training was carried out for 50 epochs using random sampling to correct for class imbalance.To improve interpretability and reduce noise, a second training step was performed after pruning the input features to retain the 100 genes with the highest weights in the first layer. The final model was validated on the held-out test set and applied to the integrated dataset to predict the identity of each single cell. Clusters were assigned a label based on the majority class prediction (>60% of cells within a cluster). Final annotations (Extended Data Fig. 1B, Extended Data Fig. 3B) were manually curated by combining the predicted identities with the expression of known marker genes.

Annotation of adult astrocyte subtypes was achieved by integrating the most specific DEG of each cluster with spatial localization predictions. Specifically, cells were assigned to one of four major astrocyte classes - Bergmann glia (BG), granular layer astrocytes (GLA), white p>matter astrocytes (WMA), or cerebellar nuclei astrocytes (CNA) - based on their predicted regional distribution within the cerebellum (i.e., localization to the PCL/ML, GL, WM, or CN, respectively). Spatial assignment was informed by three complementary sources: (i) visual inspection of identity transfer results from the Visium ST dataset; (ii) visual inspection of identity transfer on the MERFISH cerebellar slices; and (iii) results from the quantitative layer enrichment analysis. When at least two of these three lines of evidence were concordant, the corresponding regional identity was assigned to the cluster.

Annotation of astrocyte subtypes within the postnatal *Slc1a3* subset was achieved by subsetting and re-integrating (as described above) cells from the more mature astrocyte clusters (i.e. clusters 9, 14 and 20) and collected at late developmental stages (from P10 onwards). Final subtype identities were assigned using a multi-layered approach that combined: (i) expression scores of differentially expressed genes (DEGs) characteristic of adult astrocyte subtypes (Supplementary Table 1, Fig. 2f); ii) the expression of adult subtype-specific genes (Fig. 2g); and iii) spatial mapping onto the adult Visium mouse cerebellum ST dataset.

### Differential gene expression (DEG) analysis

Differential gene expression analysis was performed on R with the Seurat function FindAllMarkers() using Wilkoxon Rank Sum test, retaining only genes with an average log2 fold change (avgLog2FC) major or equal to 0.5 and expressed by at least 25% of the members of the cluster.

### Gene ontology (GO) analysis

GO enrichment analysis was performed using the ViSEAGO package^99^. DEGs identified from the pairwise comparison between Bergmann and Astrocyte groups were analyzed for enrichment of biological process GO terms using Fisher’s exact test. Only terms with a p-value

≤ 0.05 were considered significantly enriched. Enriched terms were then clustered based on semantic similarity, and manual curation was performed to highlight the most informative terms relevant to adult astrocyte biology. The same procedure and statistical parameters were applied independently to each astrocyte subtype.

### Pseudotime Computation

Pseudotime analysis of the *Slc1a3*-expressing postnatal subset was performed using the Monocle3 R package^100^. The Seurat object was converted into a *cell_data_set* Monocle object. Size factors were estimated, and dimensionality reduction and preprocessing were carried out prior to Leiden clustering on the UMAP embedding previously computed with Seurat using the cluster_cells() function, allowing identification of a single main partition. A trajectory graph process was then computed. Root nodes were selected based on their proximity to cells with high proliferation gene scores and belonging to perinatal clusters (i.e., clusters 0, 1, 2, 3, 4, 5, 6, 7, 10, 11, 15, 17, 18, and 21; see Fig. 2D).

### Maturational trajectories and lineage Inference

To infer the maturation trajectories within the postnatal *Slc1a3*-expressing subset, we employed three distinct algorithms to compute fate probabilities (i.e. the probability of each cell to be part of a maturation trajectory of a defined lineage). Two of these methods were implemented using the CellRank2 package^101^, while the third was based on the URD framework^102^. Given the computational nature of these analyses and the inherent complexity of developmental processes, we reasoned that combining multiple inference strategies would enhance the robustness and consistency of the resulting trajectories.

#### Anterograde random walks with CellRank2

Three distinct transition matrices (TMs) were generated in CellRank2 to model different aspects of cellular progression: one based on pseudotime values previously inferred with Monocle3 (pseudotime kernel), one based on experimental time using the postnatal age of tissue collection (experimental time kernel), and one based on transcriptional similarity among cells (connectivity kernel). These TMs were used to simulate biased anterograde random walks (aRWs), favoring developmental progression. Two separate simulations were performed: the first combined the pseudotime TM with the experimental time TM, while the second combined the pseudotime TM with the connectivity TM. In both cases, starting points were defined as cells collected at P0, belonging to the perinatal clusters (as described above), and displaying pseudotime values below 2.5. The inference of fate probabilities was carried out over multiple iterative rounds of random walk simulations. In the initial round, terminal states were predefined as mature astrocyte subtypes, oligodendrocytes, interneurons, and ependymal cells. In subsequent rounds, terminal states were redefined based on the clusters most enriched in fate probability from the previous round. Each round yielded a fate probability matrix indicating the likelihood of each cell reaching each terminal state. Probabilities were then averaged across cells belonging to the same cluster, generating an estimate for each cluster’s involvement in the maturation trajectories. Finally, these values were normalized such that, for each terminal state, the sum of probabilities across all clusters equaled 1.

#### Retrograde random walks with URD

Maturation trajectories were also inferred using the URD R package. A URD object was created from the subset of *Slc1a3*-expressing cells. A diffusion map was computed on the integrated UMAP previously generated with Seurat, using a sigma value of 20. URD pseudotime was then calculated by selecting as roots the cells collected at P0, belonging to the perinatal clusters (as described above), and displaying pseudotime values below 2.5. These same cells were also used as roots to simulate retrograde random walks (rRWs).

As with CellRank aRWs, multiple rounds of rRWs were performed. In the initial rounds, simulations were biased to start from mature astrocyte subtypes, oligodendrocytes, interneurons, or ependymal cells. In subsequent rounds, the top-ranked clusters from the previous round were used as new starting points.

Each round yielded a visitation frequency matrix indicating the likelihood that a walk initiated from a specific starting point would transition through each cell *en route* to the roots. Visitation frequencies were then averaged across within the same cluster to estimate each cluster’s involvement in the maturation trajectories. These values were finally normalized so that, for each starting point, the sum of probabilities across all clusters equaled 1.

#### Definition of consensus results

To generate a consensus estimate of fate probabilities across methods, the matrices obtained from the two CellRank2 analyses and the URD analysis were combined into a single consensus matrix.

First, a logit transformation was applied to each individual probability (*P*) that cluster *i* contributes to the terminal state *j* as computed by method *k*:

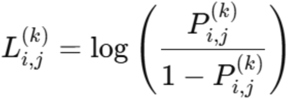

These logit-transformed values (*L*) were then averaged across methods:

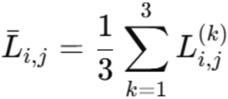

The resulting averaged values were then transformed back into probabilities using the inverse logit function:

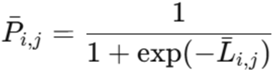

Finally, the probabilities were normalized such that for each terminal state *j*, the sum of probabilities across all clusters *i* equaled 1:

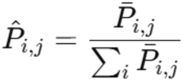

This approach yielded a robust consensus estimate of the likelihood that each cluster contributes to the maturation trajectory toward a given terminal state.

#### Validation of consensus results

To validate the predictive accuracy of our trajectory inference strategy, we leveraged the oligodendrocyte lineage, which undergoes well-characterized differentiation steps marked by stage-specific gene expression^103,104^ (Fig. 2C; Extended Data Fig. 5D–F). In the first round of random walks, the model accurately predicted that mature oligodendrocytes (Cluster 13, characterized by high *Mog* expression) originate from pre-myelinating oligodendrocytes (Cluster 16, marked by *Bcas1* expression). In the second round, Cluster 16 - now designated as the new terminal state - was traced back to *Cspg4*-expressing oligodendrocyte progenitor cells (OPCs) in Cluster 8. In the third round, Cluster 8 was further predicted to derive from Cluster 18, which expresses multiple pre-OPC markers such as *Rfx4*, *Sall3*, *Ptprz1*, and *Ncald*. This strategy also successfully reconstructed the interneuron lineage within the *Slc1a3*-expressing subset. The predicted trajectory progressed from transient amplifying progenitors (TAPs) in Cluster 11 - co-expressing *Ptf1a*, *Ascl1*, and *Mki67* - to *Lhx1/Lhx5/Mki67*+ TAPs in Cluster 12, and ultimately to *Pax2/Gad1/Gad2*+ interneuron precursors in Cluster 5^32,105,106^ (Extended Data Fig. 5G).

This approach successfully reconstructed differentiation steps up to clusters enriched at intermediate stages (Fig. 2D, middle panel). However, additional rounds of random walks to establish hierarchical relationships among clusters enriched at perinatal stages (Fig. 2D, top panel) were not feasible due to intrinsic constraints of the methods, which requires the definition of root cells and tips - two sets that cannot overlap by definition - and the assignment of a pseudotime score, which is uniformly low and similar across cells collected at the earliest stages (Extended Data Fig. 5B).

### Lineage coupling correlations analysis

To investigate lineage relationships among the clusters within the Slc1a3 postnatal subset (designated as “Slc1a3_” followed by the cluster number, e.g., “Slc1a3_cl0”) and the full postnatal dataset from which were removed cells with no neural tube origin (Microglia and Fibroblasts) and with no clear identity, we analyzed clonal overlaps, lineage coupling z-scores, and correlation patterns, adapting methodologies previously described^40,42,93^. In this analysis, a TrackerSeq clone was considered to be “shared” between two groups if it included at least two cells assigned to each cluster. We counted the number of shared clones between every pair of clusters and then compared these counts to a null distribution generated by randomly permuting the cluster annotations across 10,000 iterations. From this, we computed a z-score for each observed value to quantify the deviation from random expectations. Subsequently, we calculated Pearson correlation coefficients between the z-score profiles of all pairs to assess the similarity in their clonal sharing patterns. These correlation matrices were visualized through hierarchical clustering. For further methodological details regarding the computation of lineage coupling and correlation metrics, please refer to the original TrackerSeq publication^40^.

### Scenic gene regulatory network analysis

Single-cell regulatory network inference and cell-type–specific transcription factor (TR) activity were assessed using the pySCENIC pipeline^107^. The pipeline was executed with the command line interface of pySCENIC 0.12.1, encapsulated in a Podman container to ensure reproducible software dependencies and environment. The input consisted of the gene-expression matrix containing the counts of the subset of *Slc1a3*-expressing cells from the postnatal dataset.

First, co-expression modules were inferred with the pyscenic grn command. This step identifies sets of genes that exhibit correlated expression patterns across cells. The co-expression analysis was performed using grnboost2, which ranks potential TR–target relationships based on shared expression patterns. The output from this step is a table of TR– target associations that serve as candidates for subsequent refinement. Next, the initial TR–target associations were pruned and refined with the pyscenic ctx command, integrating both cis-regulatory sequence motif analyses and motif annotations from https://resources.aertslab.org/cistarget/. In this stage, each TR’s candidate targets are evaluated for the presence of known binding motifs in their promoter and enhancer regions. Targets lacking significant motif enrichment are removed, ensuring that putative TR–target relationships better reflect established regulatory interactions. The output from this step is a set of *regulons*, each composed of a TR and its high-confidence targets. Finally, the activity of each regulon across individual cells was quantified using the pyscenic aucell command. This calculates an “Area Under the Curve” (AUC) score that reflects the extent to which a regulon’s target genes are expressed in a particular cell.

After running the SCENIC pipeline, we linked regulon activity with the consensus matrix of maturation scores obtained as described above.For each trajectory in the consensus matrix, we computed Pearson correlation coefficients between its generation probability (consensus logit values) and each transcription factor’s AUC values. Statistical significance of these correlations was assessed using the built-in cor.test function, followed by multiple-test correction with the Benjamini–Hochberg false discovery rate (FDR). Transcription factors meeting defined significance (e.g., FDR ≤ 0.05) and minimum correlation thresholds (e.g., Pearson’s ρ > 0.1) were considered positively correlated with the corresponding lineage’s maturation trajectory.

### Immunohistochemistry

Under anesthesia, animals were transcardially perfused with an appropriate volume of 4% paraformaldehyde (PFA) in 0.12 M phosphate buffer (PB), pH 7.2–7.4. Brains were removed, stored overnight (o/n) in the same fixative at 4 °C, washed in PBS, and finally cryoprotected in 30% sucrose in 0.12 M PB. The cerebella were then embedded and frozen over dry ice in OCT (TissueTEK), sectioned in the parasagittal plane at 30 μm using a cryostat and placed directly onto glass slides (E14.5-P4 cerebella). For immunolabeling, sections were incubated o/n at room temperature with the appropriate primary antibodies dissolved in PBS with 2% bovine serum albumine (BSA, Sigma-Aldrich) and 0.25% Triton X-100 (Sigma-Aldrich): Anti-ACSA-2 (1:250, rat, monoclonal, Miltenyi Biotec), anti-Aquaporin 4 (AQP4, 1:2,500, rabbit, polyclonal, Sigma Prestige Antibodies,), anti-GAT3 (1:100, rabbit, monoclonal, Abcam), anti-GFP (1:700, chicken, polyclonal, IBA Lifesciences), anti-SOX9 (1:600, goat, polyclonal, R&D). Sections were then exposed for 2 h at room temperature to secondary species-specific antibodies conjugated with Alexa Fluor 555 (1:500; Invitrogen; anti-rabbit, anti-rat or anti-goat) or Alexa Fluor 488 (1:500, Invitrogen, anti-chicken). Cell nuclei were visualized using 4′,6-diamidino-2-phenylindole (DAPI; Fluka). After processing, sections were mounted on microscope slides with Tris-glycerol supplemented with 10% Mowiol (Calbiochem).Histological specimens were examined using a Nikon E-800 microscope, and high-resolution images were acquired with the Zeiss Axioscan slide scanner.

## Acknowledgments

We thank professor Paola Berchialla for constructive help with the analyses.

## Author contributions

V.C., A.B. and L.T. conceived the project and designed the experiments; V.C., I.V., B.X., L.S.-F, A.L., and E.M. performed the experiments; R.B., M.G, J.F.-S. contributed data, conceptualization; E.B., contributed methodology; V.C., G.T. and L.T. analyzed the data; V.C., G.T., A.B. and L.T. wrote the manuscript with input from all authors.

## Funding

This work was funded by the ERC starting grant (to L.T.; CERDEV_759112), the SNSF grant (to L.T.; 31003A_182676/1), the European Union’s funded Consortium NSC-Reconstruct: Novel Strategies for Cell based Neural Reconstruction (to A.B.; H2020, GA no. 875758), the Banca d’Italia grant (to A.B. and V.C.; Spacer, 2022), the European Advanced infraStructure for Innovative Genomics, H2020 EASI-Genomics grant PID15185 (to A.B. and V.C.), the Italian Ministry of Research and Education (MIUR) national project “Dipartimenti di Eccellenza 2018–2022 and 2023-2027” awarded to the Department of Neuroscience “Rita Levi Montalcini” (University of Turin). Valentina Cerrato was supported by a Post-doctoral fellowship by Fondazione Umberto Veronesi (“Post-doctoral Fellowship Travel Grant-anno 2020” to V.C.), by salaries on the Italian MUR grant PRIN-PNRR 2022 (ID: P20225Z3J5 to E.B.) and on the NSC-Reconstruct grant.

## Extended Data Figures

**Extended Data Fig. 1.**
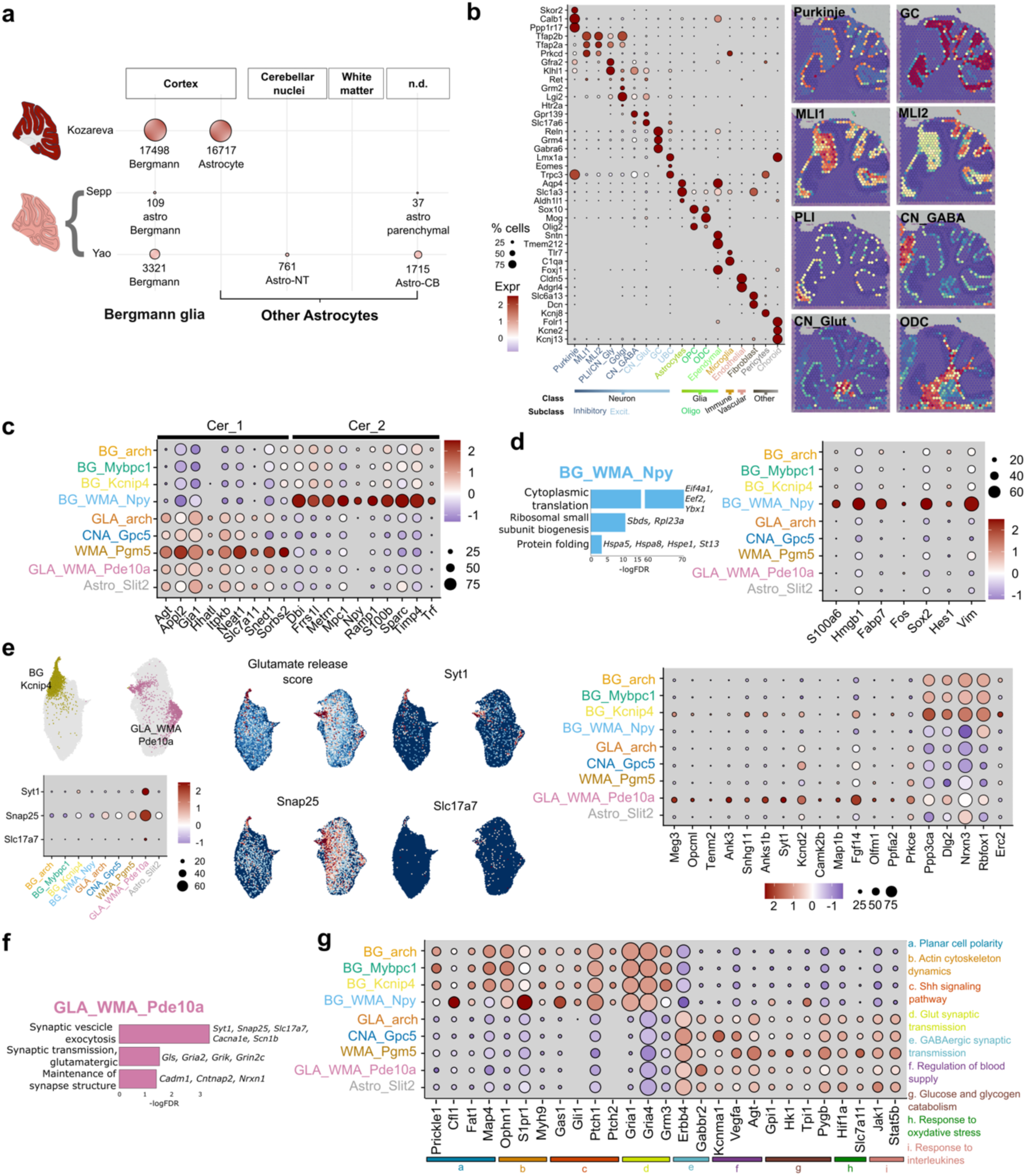
Molecular and functional annotation of the distinct clusters. a, Schematic representation of the astrocyte sampling in publicly available datasets shows the low enrichment of non-cortical astrocytes. Circle size is proportionate to the number of cells sampled. Names correspond to the annotations used by the authors of the datasets. **b,** Expression of marker genes used for cell type annotation.The bottom panel shows the classification of cell types into classes (Neurons, Glia, Immune, Vascular, Fibroblast, Pericytes, Choroid) and subclasses (Ihhibitory and Excitatory neurons, Astrocytes, Oligodendrocytes, Ependymal cells, Immune, Vascular, Fibroblast, Pericytes, Choroid). Insets on the right show the spatial mapping of selected cell types in the Visium dataset. **c,** Expression of selected DEGs of WM astrocytes belonging to the Cer_1 and Cer_2 clusters in Bocchi et al.^22^ highlights correspondence with the WMA_*Pgm5* or BG_WMA_*Npy* respectively. **d,** GO analysis (left) and gene expression (right) point to a NSC-like signature of BG_WMA_*Npy* cells. **e,** Expression of *Syt1*, *Snap25* and *Slc17a7* - either individually or combined in a score - show an enrichment in the GLA_WMA_*Pde10a* and BG_*Kcnip4* clusters. The dotplot on the right shows the enrichment of selected DEGs of hippocampal “glutamatergic” astrocytes^29^ in the same clusters. **f,** GO analysis performed on the DEGs of the GLA_WMA_*Pde10a* cluster shows an enrichment of terms associated with synaptic transmission and vesicular exocytosis. **g,** Expression of genes associated with selected enriched GO terms in Bergmann vs Astro.

**Extended Data Fig. 2.**
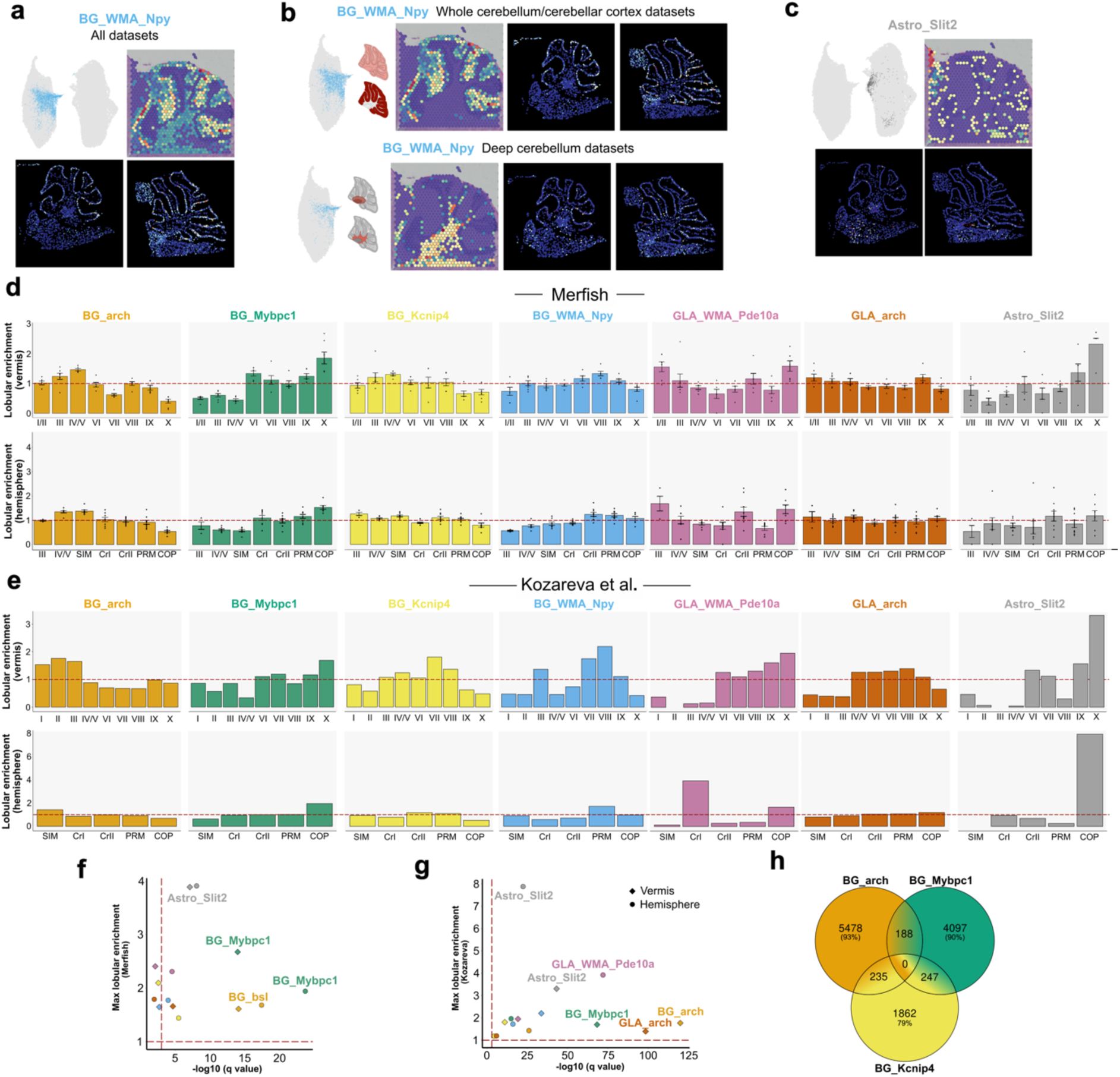
Mapping adult cerebellar astrocyte clusters through ST. a, Spatial mapping of BG_WMA_*Npy* (highlighted in the UMAP plot, top left) in the Visium (top right) or Merfish (bottom) datasets. **b,** Spatial mapping of the cells belonging to the BG_WMA_*Npy* cluster sampled from the whole cerebellum and cerebellar cortex (Yao^20^ and Kozareva^18^ datasets, top) or from the deep cerebellum (our and Bocchi^22^ datasets, bottom) highlights an enrichment in the ML/PCL or WM, respectively. The bar plots show the average layer enrichment calculated from Merfish predictions. **c,** Spatial mapping of Astro_*Slit2* in the Visium or Merfish^20^ datasets show a broad distribution of these cells across all regions. **d,** Lobule enrichment in the vermis (top) or hemisphere (bottom) calculated from Merfish predictions for the clusters that populate the cortical layers. Bars indicate the average enrichment across multiple sections 土 SEM (n=7, vermis; n=9, hemisphere). **e,** Lobule enrichment in the vermis (top) or hemisphere (bottom) calculated for the astrocyte clusters that populate the cortical layers based on tissue sampling of the snRNA-seq Kozareva18. The spatial distribution of the BG_Mybpc1 cluster is consistent with that of the Mybpc1-expressing Bergmann_2 cluster in the same study. **f,g,** Clusters that show a lobule enrichment pattern in Merfish predictions (**f**) or Kozareva18 dataset (**g**) which is significantly different from the cerebellar astrocyte population as a whole (Pearson’s chi-squared); x axis shows −log10-transformed q values, with q value < 0.001 indicated by dashed vertical line; y axis shows the maximum lobule enrichment for each type/subtype, with lobule enrichment = 1 (comparable to a random distribution) indicated by dashed horizontal line. **h,** Venn diagram showing the intersections across the cells localized in the ML/PCL predicted as BG_*arch*, BG_*Mybpc1* or BG_*Kcnip4* in the Merfish^20^ dataset highlights a low level of overlap across the three populations (n= predicted cells).

**Extended Data Fig. 3.**
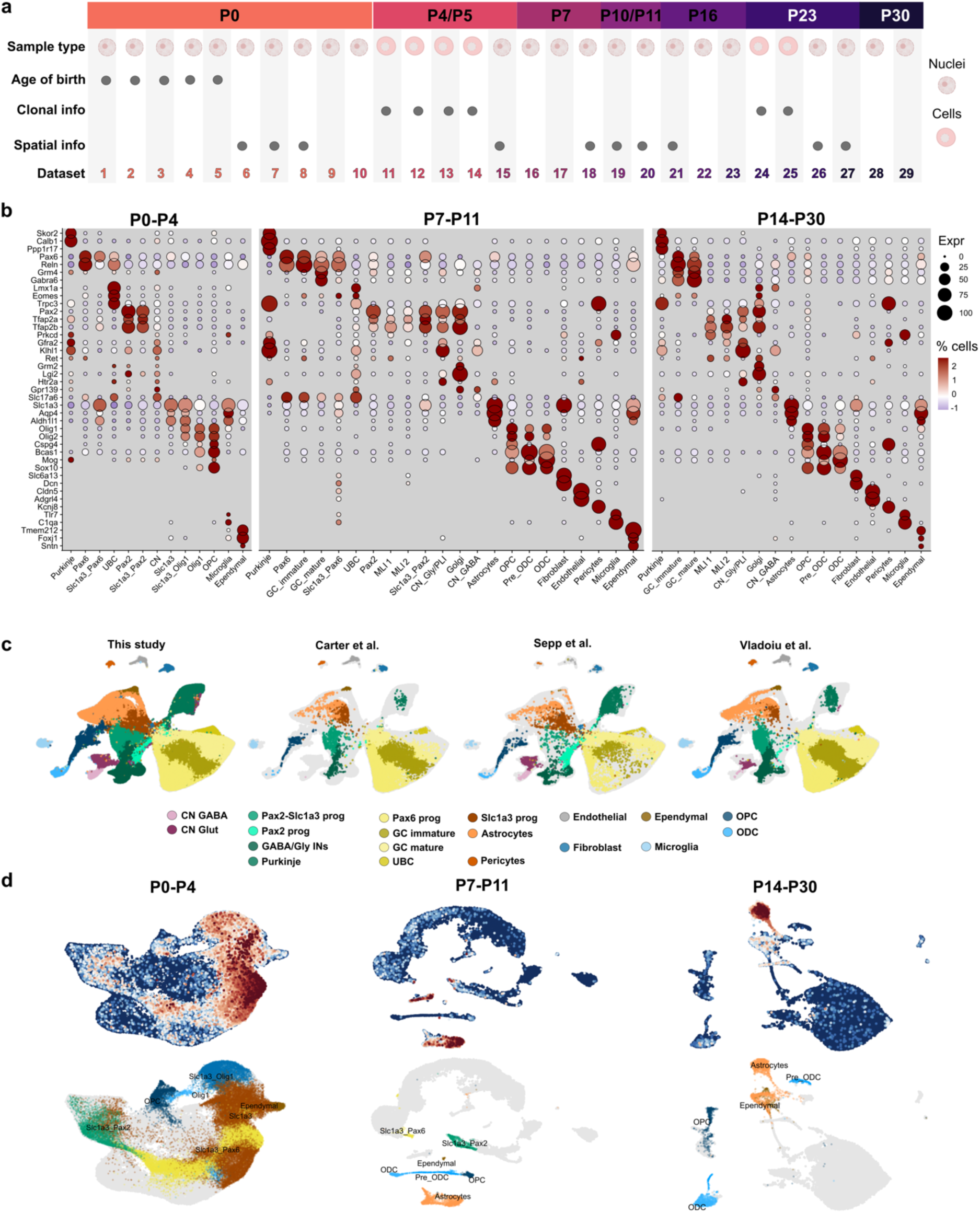
Overview of the multi-modal sc/snRNA-seq dataset of the whole postnatal mouse cerebellum. a, Schematic representation of the information contained within each experimental set of the newly generated postnatal dataset, sampling nuclei or cells across several developmental stages from P0 to P30. Biological and technical replicates are not shown. **b,** Expression levels of key cell type-specific markers used for cell annotation. **c,** Comparison of our postnatal dataset with publicly available datasets^19,32,33^, highlighting the improved coverage of the full repertoire of cerebellar cell lineages. UMAP plots display the integrated dataset across all developmental stages, split by dataset and colored by cell type. **d,** UMAP plots of the P0-P4, P7-P11 or P14-P30 integrated newly generated datasets showing *Slc1a3* expression (top panels) and the *Slc1a3*-expressing clusters that were subsequently subset for further analyses (bottom panels).

**Extended Data Fig. 4.**
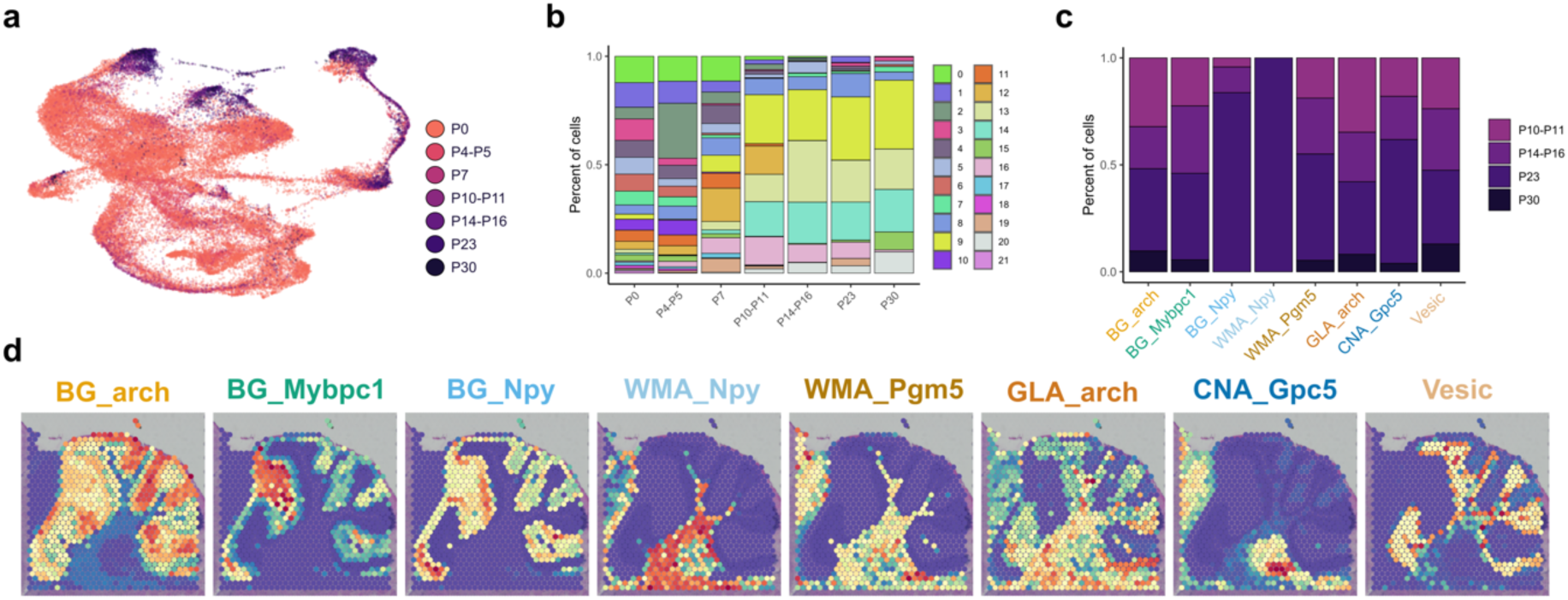
Temporal dynamics and profiling of Slc1a3-expressing cells during postnatal cerebellar development. a, UMAP plot of the postnatal subset, colored by stage of sampling. **b,** Barplot showing cluster distribution across distinct stages. Clusters 9, 14 and 20, which contain differentiated astrocytes, become predominant after P10-P11. **c,** Bar plot illustrating the fraction of cells collected at each developmental stage after P10-P11 for each annotated astrocyte subtype. **d,** Spatial mapping of the astrocyte subtypes identified within the postnatal subset onto the adult mouse sagittal section of the Visium dataset.

**Extended Data Fig. 5.**
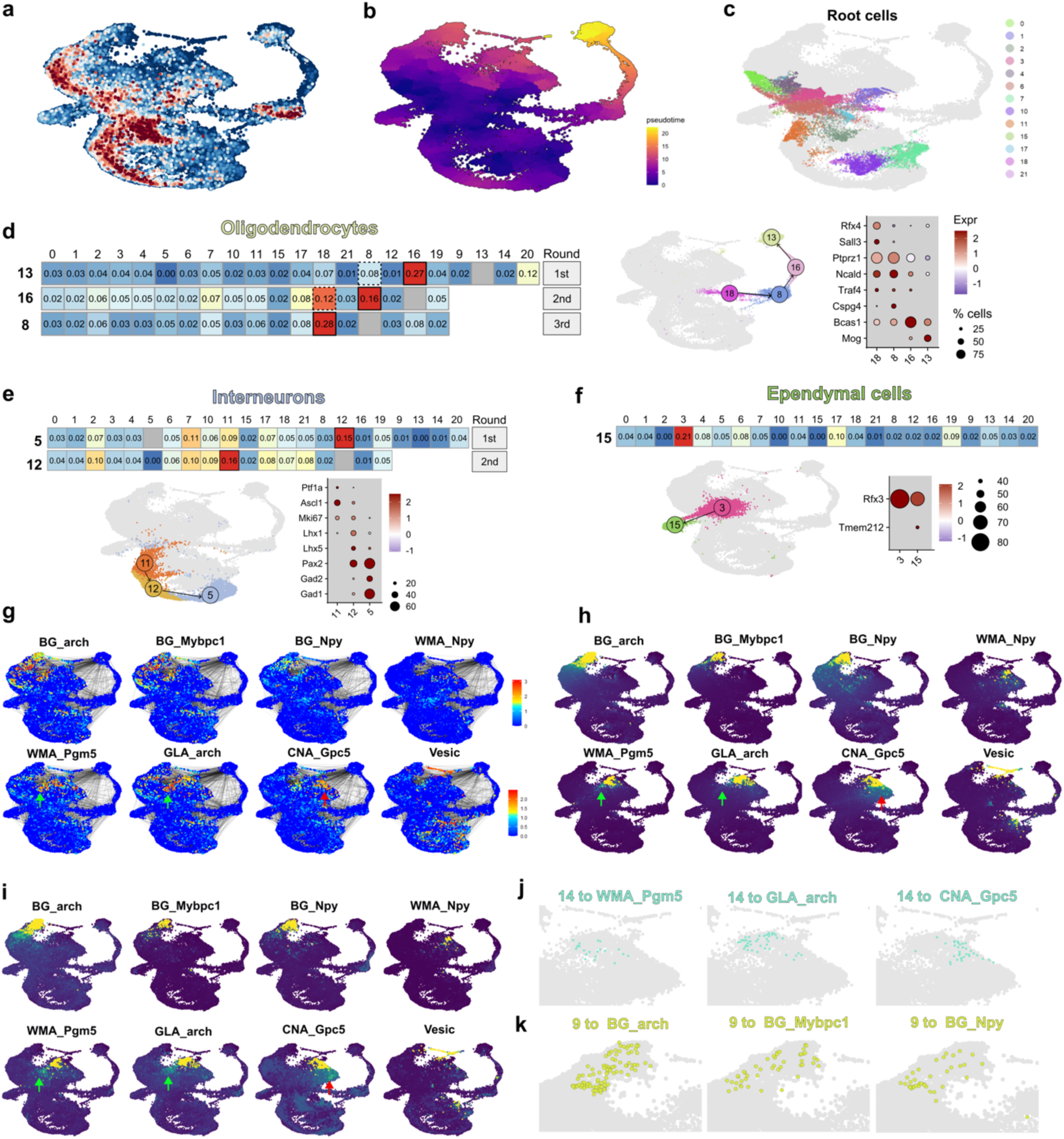
Details and validation of the combined computational approach to predict the maturation trajectories. a, UMAP plot of the postnatal subset of *Slc1a3*-expressing cells, with cells colored according to their computed proliferation score. This score was used as a starting point to calculate the pseudotime using Monocle3^100^ (**b,** see Methods). **c,** UMAP highlighting the cells used as roots for random walk simulations, colored by cluster. **d,** Validation of the predictive power of the approach using the oligodendrocyte lineage (see Methods). **e,** Validation of the predictive power of the approach using the interneuron lineage (see Methods). **f,** Ependymal cells, corresponding to cluster 15, were predicted to originate from cluster 3, both of which consistently exhibit high levels of *Rfx3*, a key TR required for ependymal cell differentiation^108^. **g-i**, UMAP plots showing visitation probability values obtained in the first round of random walks simulated with URD (**g**) or CellRank (**h,i**) from or toward each of the mature astrocyte subtypes defined as tips. Green arrows point to cells in cluster 14 with a high probability of differentiating into WMA_*Pgm5* or GLA_*arch*. Red arrows indicate cells in cluster 14 with a high probability of differentiating into CNA_*Gpc5.* This is consistent with the results obtained in the additional round of random walks shown in Fig. 3c. **j,k,** UMAP insets highlighting the cells in cluster 14 (j) or 9 (k) with the highest probability of differentiating into WMA_*Pgm5*, GLA_*arch*, CNA_*Gpc5* or BG_*arch*, BG_*Mybpc1*, BG_*Npy*, respectively. Notably, cells in cluster 14 show minimal overlap, indicating distinct pools of committed precursors for WMA_Pgm5, GLA_*arch*, and CNA_Gpc5.

**Extended Data Fig. 6.**
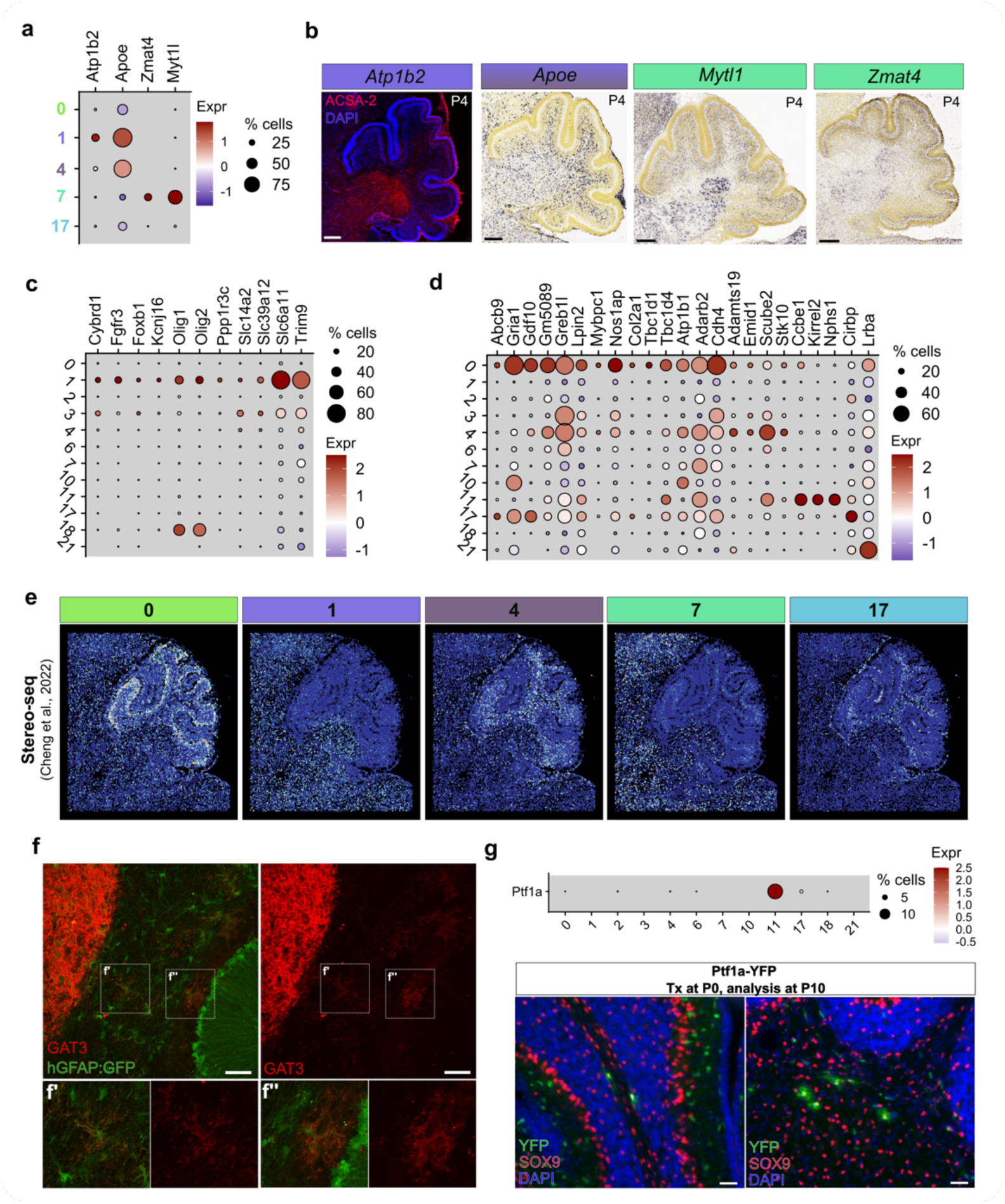
Profiling of perinatal progenitor clusters. a, Expression levels of key genes differentially expressed in cluster 1, cluster 4 or cluster 7 that also show distinct spatial expression patterns as indicated by ISH data (Allen Brain Atlas) or immunofluorescence (**b**). **c,** Genes enriched in the ACSA2/GLAST+ progenitor pool^16^ are also highly expressed in cluster 1. **d,** Genes differentially expressed by the ACSA2-/GLAST+ progenitor pool^16^ show low expression in cluster 1 but high expression in other clusters of the subset, namely cluster 0, 4 and 11, consistent with the multipotency of this progenitor pool along both interneuron and glial lineages. **e,** Spatial mapping of the perinatal astrocyte progenitor clusters onto a P7 Stereo-seq dataset^37^. Although the cerebellum at P7 likely hosts few remaining cells belonging to these perinatal clusters (mostly composed of cells collected between P0 and P5, Extended data Fig. 4a), the spatial predictions align well with their inferred maturation trajectories. Cluster 0 predominantly maps to the BG layer, cluster 1 to the CN, cluster 4 to the WM and GL, cluster 17 to the WM. The predicted localization of cluster 7 within the CN and outer GL, regions populated by emerging glutamatergic neurons, is consistent with the expression pattern of its DEGs (**b**) and likely reflects their transcriptional similarity with developing excitatory cells. **f,** Immunofluorescence for GAT3 (red) in hGFAP:GFP mice at P30 highlights its expression in the CN and in few astrocytes of the deep WM (**f’**) and GL (**f’’**), further confirming the inferred fate potency of cluster 1. **g,** *Ptf1a* expression is restricted to cluster 11 early postnatally (top panel). Genetic fate mapping performed at P10 after Tx administration at P0 in Ptf1a^CreERTM^ × R26R^YFP^ mice (Ptf1a-YFP) shows that Ptf1a-expressing progenitors at P0 give rise mostly to ML interneurons and to few astrocytes localized in the WM. Scale bars: 50 µm.

**Extended Data Fig. 7.**
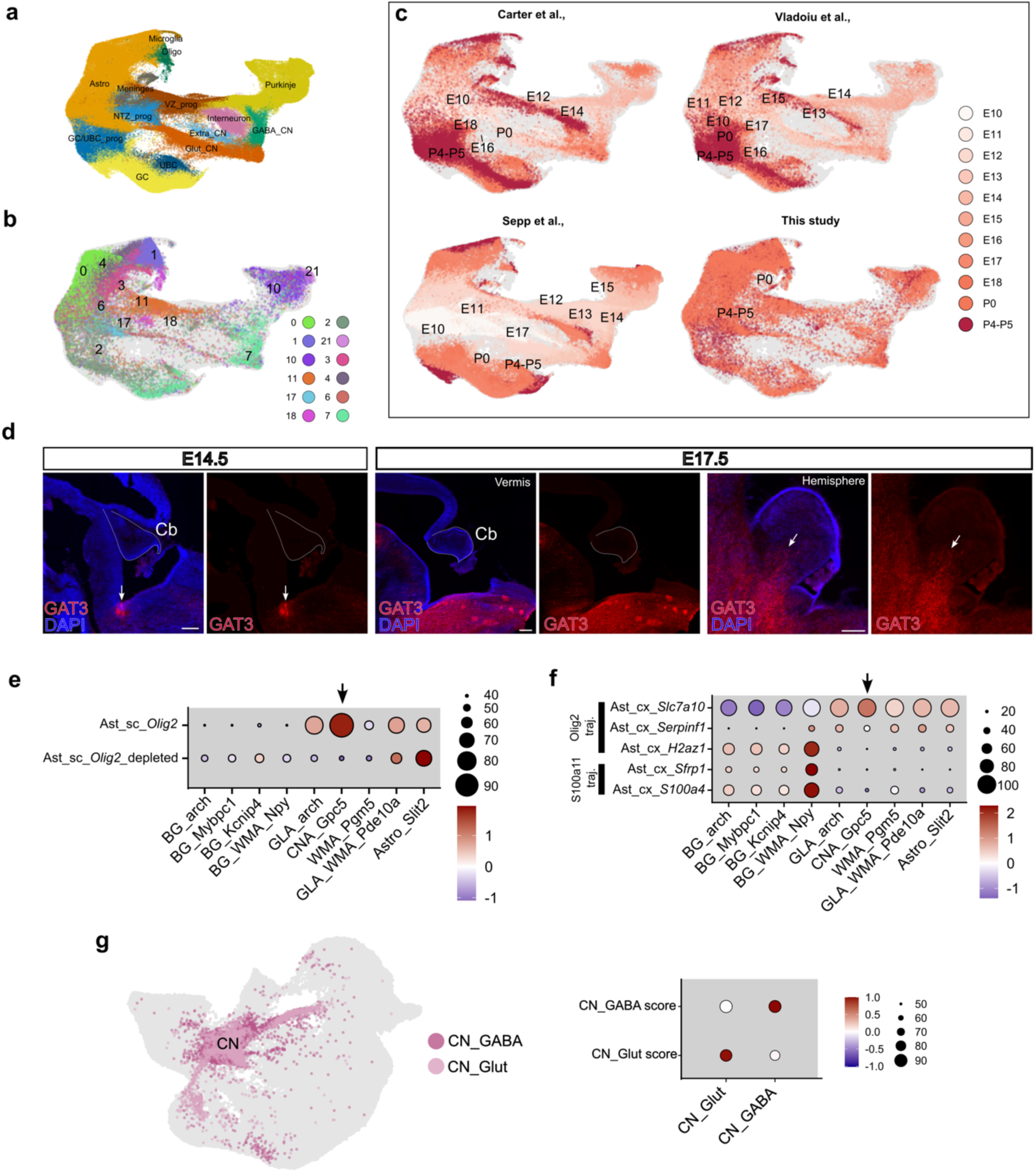
Spatial and transcriptional evidence for diverse embryonic origins of cerebellar astrocytes. a, UMAP of the dataset obtained by integrating our *Slc1a3*-expressing perinatal clusters (Figure 2D, top panel) and publicly available embryonic and early postnatal cerebellar sc/snRNA-seq datasets^19,32,33^, with dots colored by cell type/lineage. **b,** UMAP highlighting, within the integrated dataset, the cells belonging to the *Slc1a3*-expressing clusters from our dataset, colored by cluster identity. **c,** UMAP plots of the integrated dataset, split by study and colored by developmental stage. **d,** Immunofluorescence for GAT3 (red) at E14.5 or E17.5, showing its initial high expression in the pontine hindbrain at E14.5, followed by broader hindbrain expression and selective cerebellar localization in the CN region by E17.5; note the absence of signal in the vermal cerebellum at E17.5. **e,** Dot plots showing expression scores of DEGs from spinal cord Olig2-expressing (Ast_sc_Olig2) and Olig2-negative (Ast_sc_Olig2_depleted) astrocyte populations^43^ across the adult cerebellar astrocyte subtypes. **f,** Dot plots displaying the expression scores of DEGs from cortical astrocyte clusters derived from either Olig2-progenitor or S100a1-progenitor lineages^42^ across the adult cerebellar astrocyte subtypes. **g,** UMAP highlighting developing CN neurons within the integrated P0-P4 dataset, annotated as GABAergic (CN_GABA) or glutamatergic (CN_Glut) based on the expression scores of DEG signatures identified in the corresponding adult CN neuron populations (Fig. 1a).

**Extended Data Fig. 8.**
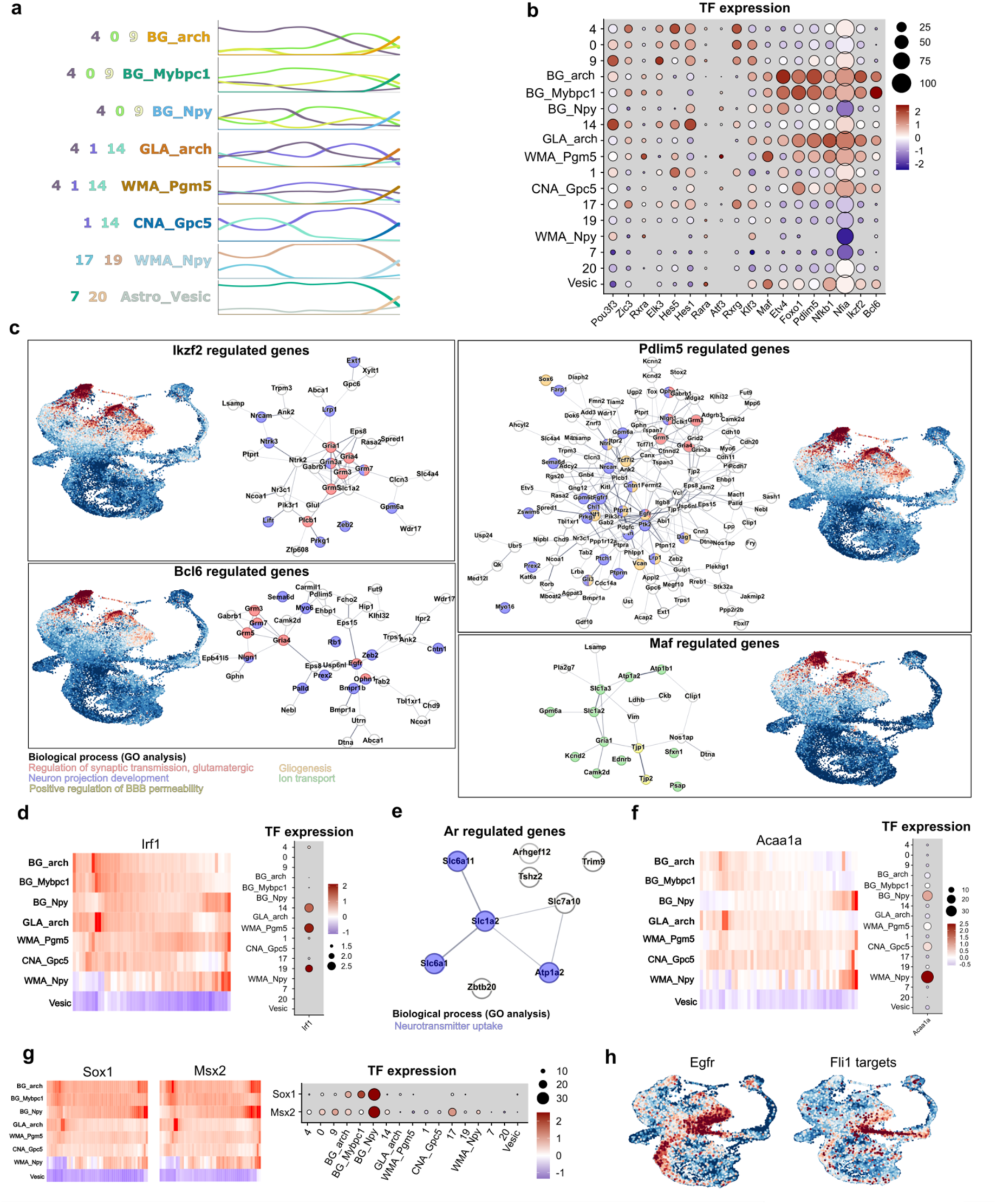
Additional characterization of SCENIC-inferred TRs. a, Cluster distribution along maturation trajectories. Lines represent the percentage of cells belonging to distinct clusters across bins (total of 50 bins per trajectory, y-axis maximum = 100%). Each subpanel illustrates the cluster dynamics specific to a single maturation trajectory. Trajectories are presented in the same order as in TRs activity heatmaps. **b,** Expression of TRs whose activity is shown in Fig. 5c. **c,** Expression scores for target genes of novel candidate regulators of cerebellar astrocyte maturation. Networks in each panel show the protein-protein interactions inferred using the STRING database (http://string-db.org). Relevant biological processes (GO analysis performed on STRING) are highlighted in distinct colors. Genes lacking interactions or involvement in relevant biological processes are excluded. **d,** Predicted activity (left) and gene expression (right) of *Irf1,* enriched in WMA trajectories. **e,** STRING-inferred protein-protein interactions among Ar target genes, indicating enrichment in neurotransmitter uptake. **f,** Predicted activity (left) and gene expression (right) of *Acaa1a,* specifically enriched in Npy trajectories. **g,** Predicted activity (left) and gene expression (right) of *Sox1* and *Msx2,* enriched in BG_*Npy*. **h,** *Egfr* expression and Fli1 target genes expression score, both high in cluster 1 and 18, at the onset of the oligodendrocyte lineage.

## Notes

### Competing Interest Statement

The authors have declared no competing interest.

